# Altered cellular metabolic pathway and epithelial cell maturation induced by MYO5B defects are partially reversible by LPAR5 activation

**DOI:** 10.1101/2024.09.03.610579

**Authors:** Michael Momoh, Sudiksha Rathan-Kumar, Andreanna Burman, Monica E Brown, Francisca Adeniran, Cynthia Ramos, James R Goldenring, Joseph T Roland, Izumi Kaji

**Affiliations:** Section of Surgical Sciences and Vanderbilt University Medical Center Nashville, TN, USA; Epithelial Biology Center, Vanderbilt University Medical Center, Nashville, TN, USA; Department of Cell and Developmental Biology, Vanderbilt University, Nashville, TN, USA; Nashville VA Medical Center, Nashville, TN, USA

**Author notes:** First authors contributed equally (Michael Momoh, Sudiksha Rathan-Kumar). Address corresponding to: Izumi Kaji, PhD. Epithelial Biology Center Vanderbilt University Medical Center 2213 Garland Ave MRB4-10465H Nashville TN 37232. **Data availability**: RNA-sequencing data on GEO (https://www.ncbi.nlm.nih.gov/geo/) with Accession No. GSE260706. Summary tables of the RNA-seq data and supplementary figures are available on NIH Figshare repository (10.6084/m9.figshare.26384263).

**Keywords:** Microvillus inclusion disease, Lysophosphatidic acid receptor, Mitochondria, Mouse model, Enteroid

## Abstract

Functional loss of the motor protein, Myosin Vb (MYO5B), induces various defects in intestinal epithelial function and causes a congenital diarrheal disorder, microvillus inclusion disease (MVID). Utilizing the MVID model mice, *Vil1-Cre^ERT2^;Myo5b^flox/flox^* (MYO5BΔIEC) and *Vil1-Cre^ERT2^;Myo5b^flox/G519R^*(MYO5B(G519R)), we previously reported that functional MYO5B loss disrupts progenitor cell differentiation and enterocyte maturation that result in villus blunting and deadly malabsorption symptoms. In this study, we determined that both absence and a point mutation of MYO5B impair lipid metabolism and alter mitochondrial structure, which may underlie the progenitor cell malfunction observed in MVID intestine. Along with a decrease in fatty acid oxidation, the lipogenesis pathway was enhanced in the MYO5BΔIEC small intestine. Consistent with these observations *in vivo*, RNA-sequencing of enteroids generated from two MVID mouse strains showed similar downregulation of energy metabolic enzymes, including mitochondrial oxidative phosphorylation genes. In our previous studies, lysophosphatidic acid (LPA) signaling ameliorates epithelial cell defects in MYO5BΔIEC tissues and enteroids. The present study demonstrates that the highly soluble LPAR5-preferred agonist, Compound-1, improved sodium transporter localization and absorptive function, and tuft cell differentiation in patient-modeled MVID animals that carry independent mutations in MYO5B. Body weight loss in male MYO5B(G519R) mice was ameliorated by Compound-1. These observations suggest that Compound-1 treatment has a trophic effect on intestine with MYO5B functional loss through epithelial cell-autonomous pathways that may improve the differentiation of progenitor cells and the maturation of enterocytes. Targeting LPAR5 may represent an effective therapeutic approach for treatment of MVID symptoms induced by different point mutations in MYO5B.

**NEW & NOTEWOTHY:** This study demonstrates the importance of MYO5B for cellular lipid metabolism and mitochondria in intestinal epithelial cells, a previously unexplored function of MYO5B. Alterations in cellular metabolism may underlie the progenitor cell malfunction observed in microvillus inclusion disease (MVID). To examine the therapeutic potential of progenitor-targeted treatments, the effects of LPAR5-preferred agonist, Compound-1, was investigated utilizing several MVID model mice and enteroids. Our observations suggests that Compound-1 may provide a therapeutic approach for treating MVID.

**Graphic Abstract:** 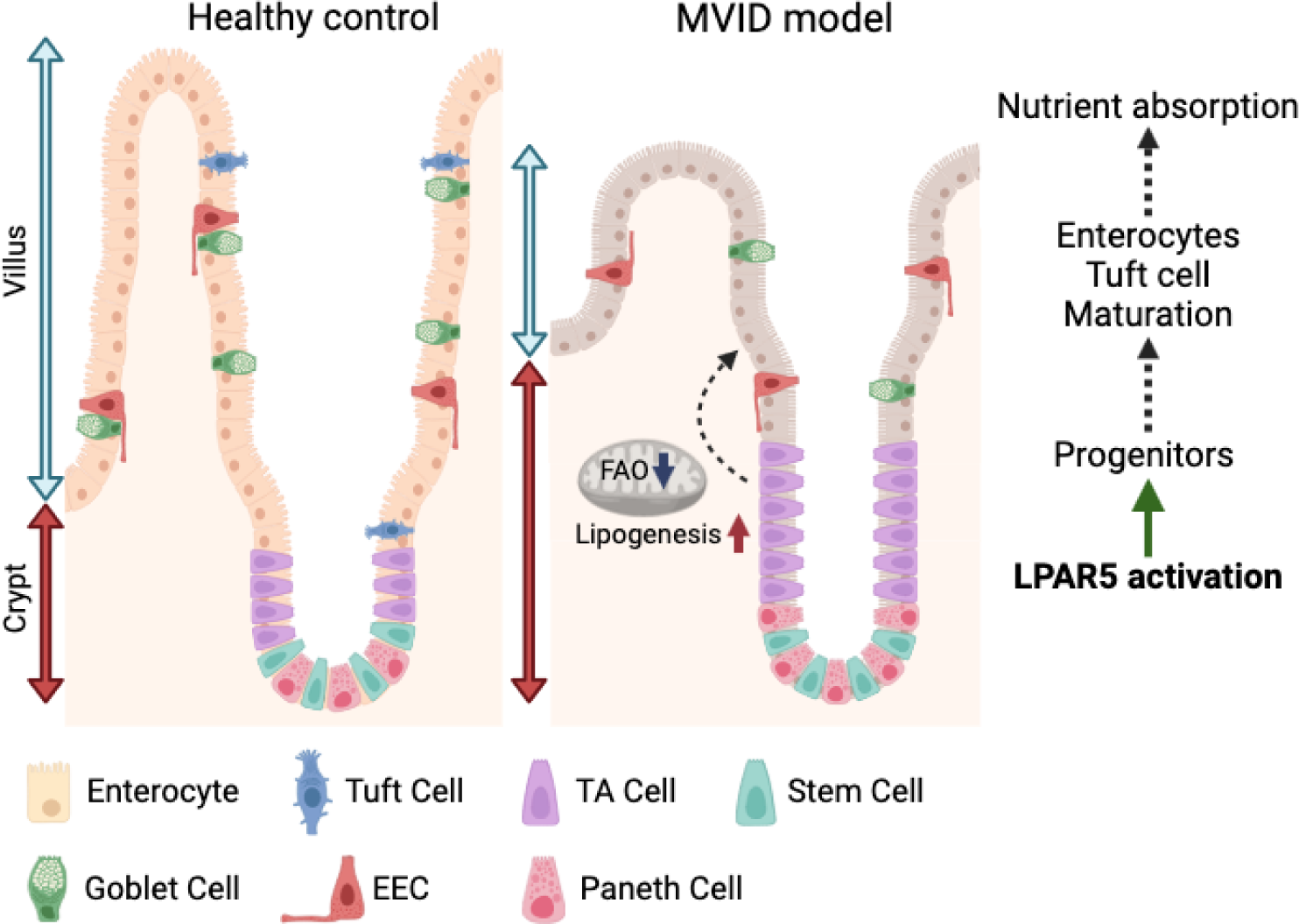

## Introduction

Microvillus inclusion disease (MVID) is an autosomal recessive disorder characterized by feeding-induced dehydrating diarrhea typically presenting in the first week after birth (1, 2). MVID is caused by homozygous or compound heterozygous inactivating mutations in Myosin Vb (MYO5B), an essential motor protein in the intestinal epithelial cells (3–5). Diarrhea in MVID patients is associated with villus blunting, loss of microvilli on the apical membrane of enterocytes, the expansion of autophagic lysosomes, and loss of apical transporters, such as sodium/proton exchanger 3 (NHE3), sodium-dependent glucose transporter 1 (SGLT1), and peptide transporter 1 (6, 7). Continuous total parenteral nutrition or intestinal transplantation are currently the only treatment options for the severe malabsorption syndrome in MVID patients (8).

To understand better the pathophysiology of intestine with inactivated MYO5B, we have established several experimental models of MVID: MYO5B deficient mouse strains (9), MVID patient-modeled mice with a compound heterozygous mutation at MYO5B(G519R) (7), and a genetically engineered pig model with a homozygous MYO5B(P663L) mutation that is homologous with the MYO5B(P660L) mutation identified in Navajo MVID patients (10). The mutant MYO5B(P660L) protein is a rigor mutant, which shows constitutive interaction with actin filament (8). On the other hand, mutant MYO5B(G519R) protein colocalizes with RAB11A and is accumulated in abnormal vesicles in subapical spaces of enterocytes, indicating a different motor status than MYO5B(P660L) (7). Intestinal tissues of these animal models commonly show expanded proliferative crypts, blunted villi, and immature microvilli structure that lacks proper localization of nutrient transporters, together phenocopying MVID patient tissues. Functionally, their expanded crypt cells maintain CFTR-mediated chloride secretion, and immature villus cells lack sodium absorption through SGLT1, all underlying MVID’s characteristic malabsorption and deadly diarrhea (11). Both inhibiting abnormal secretion and enhancing absorptive functions are critical goals for the treatment of chronic diarrhea and malabsorption. Those animal models are useful tools to investigate the altered cell biology in MVID, which can provide insights to identify additional treatment options.

We recently found that a bioactive phospholipid species, lysophosphatidic acid (LPA), can promote proper microvilli structure, suppress CFTR activity, and improve SGLT1-mediated absorption in MYO5B deficient mice (12). The effect of LPA treatment on CFTR is recapitulated by an LPA receptor (LPAR)2 selective agonist. However, the LPAR2 activation does not promote sodium absorption, indicating that other LPA receptor(s) contribute therapeutic effects on MYO5B-deficient epithelial cells (12). Further study demonstrated that LPA(18:1) treatment restores the differentiation of the tuft cell lineage in the MYO5B-deficient intestine, correlating with the maturation of enterocytes that possess proper brush border structure with sodium transporters (13). These findings suggest that the trafficking and differentiation blockades induced by MYO5B inactivation can be overcome by LPA-activated pathways, leading to the possible development of drug treatments for MVID that could obviate the need for multi-organ transplantation or long-term total parenteral nutrition.

The RNA expression of G protein-coupled receptors for LPA (LPAR) RNAs, *Lpar1*, *Lpar5*, and *Lpar6* has been identified in intestinal epithelial cells, although the topology of each receptor has not been clarified (14, 15). Previous studies have demonstrated that LPAR5 activation stimulates the trafficking of NHE3 to the apical membrane in enterocytes, which facilitates sodium and water absorption in mice (16, 17). Recent studies utilizing mouse enteroids suggest that LPARs activation is important for epithelial proliferation. LPA supplementation to the medium enhances the proliferation and differentiation of enteroids through LPAR1, independent of epidermal growth factor (18). Genetically engineered mouse enteroids also demonstrated that LPAR2 and LPAR5 reciprocally support cell proliferation (19). Furthermore, genetic deletion or chemical inhibition of LPAR5 in mouse enteroid cultures revealed that LPAR5 signaling in epithelial and non-epithelial cells is important for intestinal stem cell proliferation (20).

Based on these previous studies, we hypothesize that LPAR5 mediates the therapeutic effects of LPA treatment observed in MYO5B-deficient mice and enteroid models. Additionally, natural LPA is not an ideal treatment to obtain consistent dosing because of its poor solubility and stability. Several LPAR subtype-specific agonists and antagonists have been developed for treatment of different diseases (21, 22); however, LPAR5 agonists have not been examined in diarrheal disease models. Furthermore, the therapeutic effect of LPA signaling has not been examined on other MVID models than MYO5B-deficient mice. To test our hypothesis, we have synthesized UCM-05194 (23) and Compound-1 (24), which target LPAR1 and LPAR5, respectively. Here, we show that functional MYO5B loss impairs mitochondria and cellular metabolic pathways, which may affect epithelial cell development and that the synthetic LPAR5 agonist, Compound-1, partially rescued enterocyte maturation and tuft cell differentiation in two MVID mouse strains *in vivo* and in Navajo MVID patient-modeled pig enteroids.

## Materials and Methods

### Mice

All animal studies were performed with approval from the Institutional Animal Care and Use Committee of Vanderbilt University Medical Center (M2000104). At the 8–10 weeks of age, MYO5BΔIEC and MYO5B(G519R) mice and littermate controls (*Vil1-Cre^ERT2^;Myo5b^+/flox^*, *Myo5b^G519R/flox^*, and *Myo5b^flox/flox^*) received a single dose of tamoxifen (100 mg/kg body weight) by intraperitoneal (IP) injection (day 0) (9). Twenty-three male mice and thirty-two female mice were used in this study. LPA (18:1 1-oleoyl-Lyso PA) and GRI977143 solutions were prepared as previously reported (12), and Compound-1 and UCM-05194 were dissolved in PBS and Captisol® solution, respectively. One of the LPAR agonists or vehicle was administered by IP injection once a day, and body weight changes and diarrhea symptoms were monitored. On day 4, mice were euthanized and the duodenum (0–8 cm from the pyloric ring), jejunum (8 cm following the duodenum), ileum (distal 8 cm from the ileocecal junction), and colon were collected for tissue assessments. To investigate starvation phenotype of small intestine, four control mice were underwent food restriction for 48 hours with free access to water. The fasted mice were group-housed in regular cages with bedding. To prevent the coprophagy, cages with bedding were replaced every 12 hours.

### Imaging Mass Spectrometry

Small segments of jejunum were isolated from control and MYO5BΔIEC mice and immediately frozen in liquid nitrogen. The frozen tissues were stored at –80°C until further procedure at the Vanderbilt Imaging Mass Spectrometry Core. Tissues were sectioned at 12 μm and Matrix (9AA) was applied via TM Sprayer. Metabolites were imaged on the 15T FT-ICR at 50-μm resolution in negative ion mode. Mass spectra of molecules of interest were visualized as heatmaps.

### Electron Microscopy

After euthanasia, mouse tissues were fixed with 2.5% glutaraldehyde in 0.1 M cacodylate by trans-cardiac perfusion following warm PBS perfusion. Small pieces of intestine were further fixed in the same fixative, followed by sequential post-fixation in 1% tannic acid, 1% OsO_4_, and en bloc stained in 1% uranyl acetate. All samples were dehydrated using a graded ethanol series. TEM samples were subsequently infiltrated with Epon-812 using propylene oxide as the transition solvent, followed by polymerization at 60C for 48 hours. Samples were sectioned at a nominal thickness of 70 nm using a Leica UC7 ultramicrotome. TEM imaging was performed using a Tecnai T12 operating at 100 keV using an AMT nanosprint CMOS camera.

### Tissue immunostaining

Immunofluorescence staining and imaging were performed as previously reported (13). Four-µm-thin sections were deparaffinized and rehydrated before performing antigen retrieval using 10 mM sodium citrate buffer, contained 0.05% Tween 20 (pH 6.0), in a pressure cooker for 15 minutes. After cooling down on ice, slides were rinsed in PBS and blocked with Protein block serum-free (Dako) for 1 hour at r/t or overnight at 4°C. The primary antibodies against NDUFB8 (1:1000, Abcam ab192878), HMGCS2 (1:50, Abcam ab137043), NHE3 (1:200, Novus NBP1-82574), SGLT1 (0.5 µg/ml, own)(12), LAMP1 (1:100, Santa Cruz sc-19992), PHH3 (1:50, Novus

NBP3-08511IR, DyLight 750), Ki67 (1:100, Cell Signaling 11882S, Alexa Fluor 488), DCLK1 (1:2000, Abcam ab202755 Alexa Fluor 647), PCNA (1:50, Santa Cruz sc-56 Alexa Fluor 647), ß-catenin (12F7) (1:100, Novus NBP1-54467R, DyLight 550), villin (1:50, Santa Cruz sc-58897 Alexa Fluor 488), MUC13 (1:100, Santa Cruz, sc-390115 Alexa Fluor 546), and ACTG1 (1:100, Santa Cruz sc-65638 Alexa Fluor 546 or Alexa Fluor 790) were diluted in Dako antibody diluent with background reducing compound (S3022) and incubated on sections for 1 hour at r/t or overnight at 4°C. After rinsing in PBS, the slides were incubated with the corresponding secondary antibodies conjugated with fluorescence (Jackson Laboratory) for 1 hour at r/t. Slides were rinsed and counterstained with Hoechst 33342 (4 mM, Thermo Fisher Scientific, Waltham, MA) in PBS. Coverslips were mounted on the stained slides with ProLong Gold Antifade Reagent (P36934, Thermo Fisher). Whole slide images of antibody-labeled tissues were captured on an Aperio VERSA 200 (Leica Biosystems, Vista, CA) at the Vanderbilt Digital Histology Shared Resource (DHSR). Some immunofluorescence images were taken by using a Zeiss Axio Imager M2 with ApoTome (Carl Zeiss Microscopy, LLC, White Plains NY). Localization of NHE3 and SGLT1 in lysosomes were quantified by colocalization analysis with LAMP1 using ImageJ software (National Institutes of Health) and the plugin Coloc 2. Five representative images of jejunum per mouse were analyzed and their results were averaged.

### Digital Image Analysis

Cell segmentation and SGLT1 localization analyses have been established at VUMC DHSR on Python platform. Utilizing whole-slide fluorescence images, machine learning was performed to generate probability maps of all jejunum tissues and antibody signals, as well as tissue folds and debris (Ilastik) (25). These probability maps were used in combination with the original captured 3-4 color images to segment individual cells based on their nuclei and combined membrane marker signals using in-house coded scripts (MatLab) (26). Identified cell membranes were scored for their SGLT1 signal and positive versus negative cell cut-offs were determined automatically using an inverse slope of discretized pixel values. The ratio of membrane to cytoplasm SGLT1 intensities were calculated and converted to a heat map for easier viewing (Figure 7C). The intensities of the other studied markers: CTNNB1, Villin, and ACTG1, and their colocalization with SGLT1 were determined, as well as their membrane to cytoplasm ratios (Figure 7D: histograms). To quantify tuft cell population, DCLK1-positive cells were filtered by their size, location, and presence of a detectable nucleus. Cell number, position, and tissue area were recorded.

### Ussing chamber experiments

Mucosal-submucosal preparations were obtained from the jejunum and mounted in sliders with an aperture = 0.1 cm^2^ (Physiologic Instruments, Leno, NV) as described previously (27). Luminal and serosal surfaces of tissue were bathed in 4 ml Krebs-Ringer solution (117 mM NaCl, 4.7 mM KCl, 1.2 mM MgCl_2_, 2.5 mM CaCl_2_, 1.2 mM NaH_2_PO_4_, 25 mM NaHCO_3_, 11 Mm glucose) and maintained at 37°C using a water-recirculating heating system. Indomethacin (10 µM) was added into the serosal bath. The solution was continuously bubbled with a gas mixture of 95% O_2_ and 5% CO_2_ to maintain the pH at 7.4. Short-circuit current (*I*_sc_) was continuously recorded under voltage clamp conditions at zero potential difference by the DataQ system (Physiologic Instruments). SGLT1-dependent Na^+^ absorption was assessed by phlorizin (0.1 mM)-sensitive *I*_sc_. Cl^−^ secretion was measured by *I*_sc_ peaks after carbachol (10 µM) and forskolin (10 µM) applications to the serosal side. CFTR activity was determined using the CFTR-inhibitor, (R)-BPO-27 (10 µM). Phlorizin, forskolin (11018; Cayman Chemical), and (R)-BPO-27 (HY-19778; MedChemExpress) were dissolved in DMSO as 1000x stocks. DMSO < 0.3% in the bathing solution did not affect the *I*_sc_.

### *In vitro* induced MYO5B-knockout (iKO) and enteroids

Enteroids were generated from jejunal crypts of adult *Vil1-Cre^ERT2^; Myo5b^flox/flox^* (for *in vitro* induced KO (iKO) and control) and *Vil1-Cre^ERT2^; Myo5b^flox/G519R^* (G519R) mice without tamoxifen treatment and passaged twice with mechanical breakdown by P200 pipette. One day after the second passage, 1 μM 4-OH-tamoxifen (SML1666, Sigma) or vehicle (EtOH) was added into the Mouse Organoid Growth Medium (Stem Cell Technology) and incubated for 24 hours to induce Cre recombinase. On the next day, the medium was replaced with the differentiation medium (modified Minigut medium (28)), which contains 5% Noggin conditioned medium and 5% R-spondin conditioned medium (generated in Vanderbilt Organoid Core) to withdraw Wnt ligands and EGF for enhancing cell differentiation (13). Some induced KO (iKO) enteroids were incubated with 100 nM Compound-1 for 2 days in the differentiation conditions.

### RNA extraction and sequencing

Following the differentiation, enteroids were placed in Organoid Harvesting Solution (3700-100-01; Cultrex) for 1 hr at 4°C. Enteroids were washed with PBS and centrifuged. Enteroid pellets were immersed in TRIzol reagent (15596026; Invitrogen) supplemented with glycogen (G1767; Sigma Aldrich) and stored at –20°C. Total RNA was extracted from enteroids following the manufacturer’s instructions. RNA-sequencing with 4 samples per group and differential expressing gene analysis were performed by Novogene (Sacramento, CA).

Reference genome (Mus Musculus, GRCm38/mm10) and gene model annotation files were downloaded from genome website browser (NCBI/UCSC/Ensembl). Indexes of the reference genome was built using STAR and paired-end clean reads were aligned to the reference genome using STAR (v2.5). STAR used the method of Maximal Mappable Prefix(MMP). Alignments were parsed using the Tophat program. HTSeq v0.6.1 was used to count the read numbers mapped of each gene. And then Fragments Per Kilobase of transcript sequence per Millions base pairs sequenced (FPKM) of each gene was calculated based on the length of the gene and reads count mapped to this gene (29). Differentially expressed genes were analyzed between groups using the DESeq2 R package (2_1.6.3). The resulting *p*-values were adjusted using the Benjamini and Hochberg’s approach for controlling the False Discovery Rate(FDR). Genes with an adjusted *p*-value <0.05 found by DESeq2 were assigned as differentially expressed. Heatmaps were generated by using Heatmapper (30). This dataset of organoids is available on GEO (GSE260706). Our previously published datasets of epithelial cells, which were isolated from MYO5BΔIEC and control mouse jejunum, was analyzed as a comparison (GSE139302) (12).

### Quantitative PCR

cDNA was prepared using SuperScript III First-strand Synthesis System for RT-PCR (Thermo Fisher Scientific) from the RNA samples of epithelial cells, which were used for the previous RNA-seq (12). Real-time quantitative PCR was performed utilizing SsoAdvanced Universal SYBR Green Supermix with a CFX96 Real-Time System (Bio-Rad Laboratories) with previously published primer pairs (17, 31). Target gene expression was calculated as relative values to *GAPDH* by the ΔCt method.

### Pig MVID model enteroids

Pig MVID enteroids that possess a homozygous mutation at MYO5B(P663L) or wild type MYO5B were established from jejunal crypts as previously reported (10). Pig enteroids were expanded in Matrigel (Corning) immersed in Human Organoid Growth Medium (OGM, Stem Cell Technology). Organoid growth was assessed in 96-well plate and continuously imaged whole wells using a JuLi™ Stage (NanoEn Tek Inc., Waltham, MA). Organoid perimeter was measured by using FIJI, and organoid forming rate was calculated as sphere numbers per plated cell numbers 7 days after the passaging. *N* = 7–9 wells from each genotype. In a different set of culture, tuft cells were differentiated in Organoid Differentiation Medium (ODM, Stem Cell Technology) for 6 days, following 2 days culture in OGM after passaging. MYO5B(P663L) enteroids were incubated with 100 nM Compound-1 or vehicle (PBS) in ODM.

### Whole-mount organoid staining

Organoids were fixed with 10% NBF for 30 min in Matrigel domes, rinsed with cold PBS containing 0.1% FBS and centrifuged to remove Matrigel. Fixed organoids were blocked with 5% normal donkey serum in PBS containing 0.3% Triton X-100 for 2 hours and incubated with the primary antibodies against phosphor Girdin (1:100, IBL 28143) (32) or POU2F3 (1:200, Novus NBP1-83966) diluted with the blocking solution for 2 days at 4°C. After rinsing in PBS with 0.3% Triton X-100, donkey anti-rabbit Cy3 antibody (2.5 µg/ml), Alexa Fluor 488 conjugated phalloidin (1:400, Thermo Fisher A12379), and Hoechst 33342 (2 mM) were applied for 2 hr. Whole organoid images were taken by a Nikon Ti-E microscope with an A1R laser scanning confocal system (Nikon Instruments Inc., Melville, NY).

### Statistics

All datapoints of biological replicates were shown in graphs, and statistical differences were determined with a significant *P* value of < 0.05 using GraphPad Prism 9 statistical software. The test used in each analysis is described in the Figure legends.

## Results

### MYO5B loss negatively alters mitochondrial structures in the epithelial cells

Mitochondrial dysfunction is correlated with intestinal epithelial defects, as mitochondrial activities are involved in stem cell self-renewal and differentiation, and nutrient absorption function (33–35). Transmission electron microscopy (TEM) was utilized to investigate mitochondrial morphologies in *Vil1-Cre^ERT2^;Myo5b^flox/flox^*(MYO5BΔIEC) mouse jejunum, which demonstrates severe malabsorption compared to healthy mouse (control) tissues. As shown in **Figure 1A**, control mature enterocytes in villi possessed long, uniform microvilli measuring approximately 2 µm in length, and the subapical space contained numerous mitochondria with dense and organized cristae structures. Immature epithelial cells in crypts possessed shorter microvilli compared to villus cells and had large, circular mitochondria, suggesting that mitochondria differentially contribute to cellular metabolism in differentiated versus undifferentiated cells depending on the energy requirements of the specific cell types (36). In the MYO5BΔIEC mouse jejunum, both villus and crypt cells showed immature and disorganized microvilli and the accumulation of abnormal vesicles in the subapical space (**Figure 1A**, yellow arrows in right panel), similarly to those of the MVID patient-modeled mice, MYO5B(G519R) (7). In the MYO5BΔIEC cells of villi and crypts, mitochondria beneath the abnormal vesicles were swollen and had disorganized cristae structure, indicating a damaged and degraded phenotype. These observations suggest that a malfunction in mitochondria-mediated energy production is induced by MYO5B loss.

**Figure 1.**
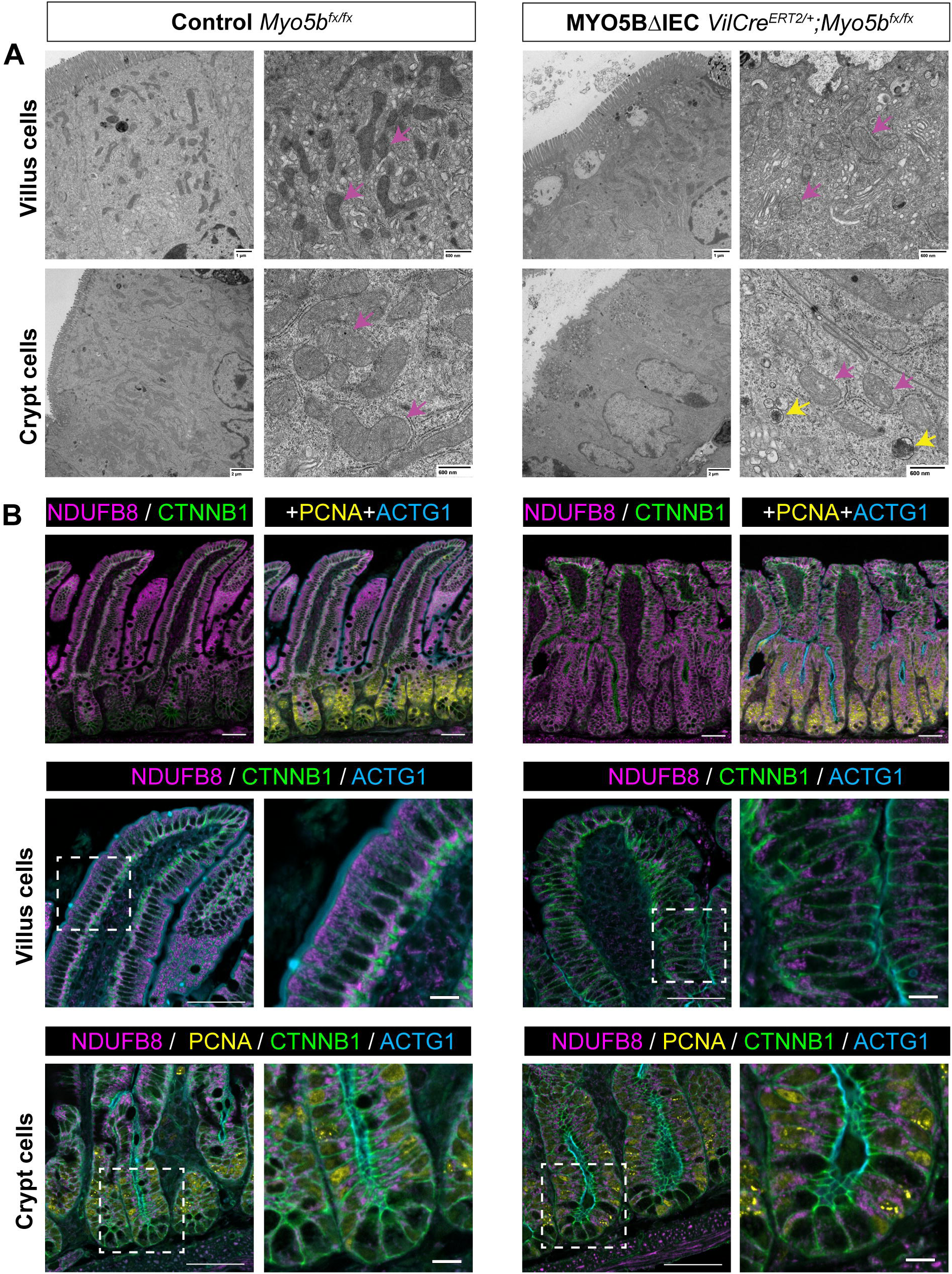
MYO5B loss negatively alters mitochondrial structures in the jejunal epithelial cells. (A) TEM micrographs demonstrate the organized microvilli and electron-dense mitochondria (pink arrows) in subapical areas of healthy control mouse intestine. In MYO5BΔIEC mouse intestine, disorganized microvilli coincide with the accumulation of subapical autophagic vesicles (yellow arrows) and swollen mitochondria (pink arrows). (B) Immunostaining for a mitochondrial complex-I protein, NDUFB8, demonstrates the sporadic mitochondrial distribution in MYO5BΔIEC mouse intestine compared to control tissues. Scale bars = 50 µm and 10 µm.

Functional mitochondria in intestinal tissues were visualized by immunostaining for a mitochondrial complex-1 protein, ubiquinone oxidoreductase subunit B8 (NDUFB8) (37) (**Figure 1B**). Control intestinal sections demonstrated dense NDUFB8 positive (+) perinuclear structures below apical actin filaments in villus enterocytes, whereas MYO5BΔIEC tissues had sporadic mitochondrial signals. The NDUFB8-negative subapical space in MYO5BΔIEC enterocytes correlated with the accumulation of autophagic vesicles in the TEM images (**Figure 1A**). PCNA+ proliferative cells in the crypts demonstrated similar cytoplasmic distributions of mitochondria to those in villus enterocytes of each mouse group.

TEM images of a healthy pediatric patient demonstrated numerous electron-dense mitochondria with organized cristae structure in the enterocytes (**Figure 2**). In the TEM sections from an MVID patient biopsy possessing the MYO5B point mutation (G519R) (7), only a few enterocytes had mitochondria in cytoplasm. As we have previously reported, this MVID patient’s biopsies demonstrated abnormal accumulation of multivesicular bodies and short microvilli. The mitochondria in the MVID patient biopsy showed swollen and disorganized cristae structure, consistent with the mouse models (**Figure 2**).

**Figure 2.**
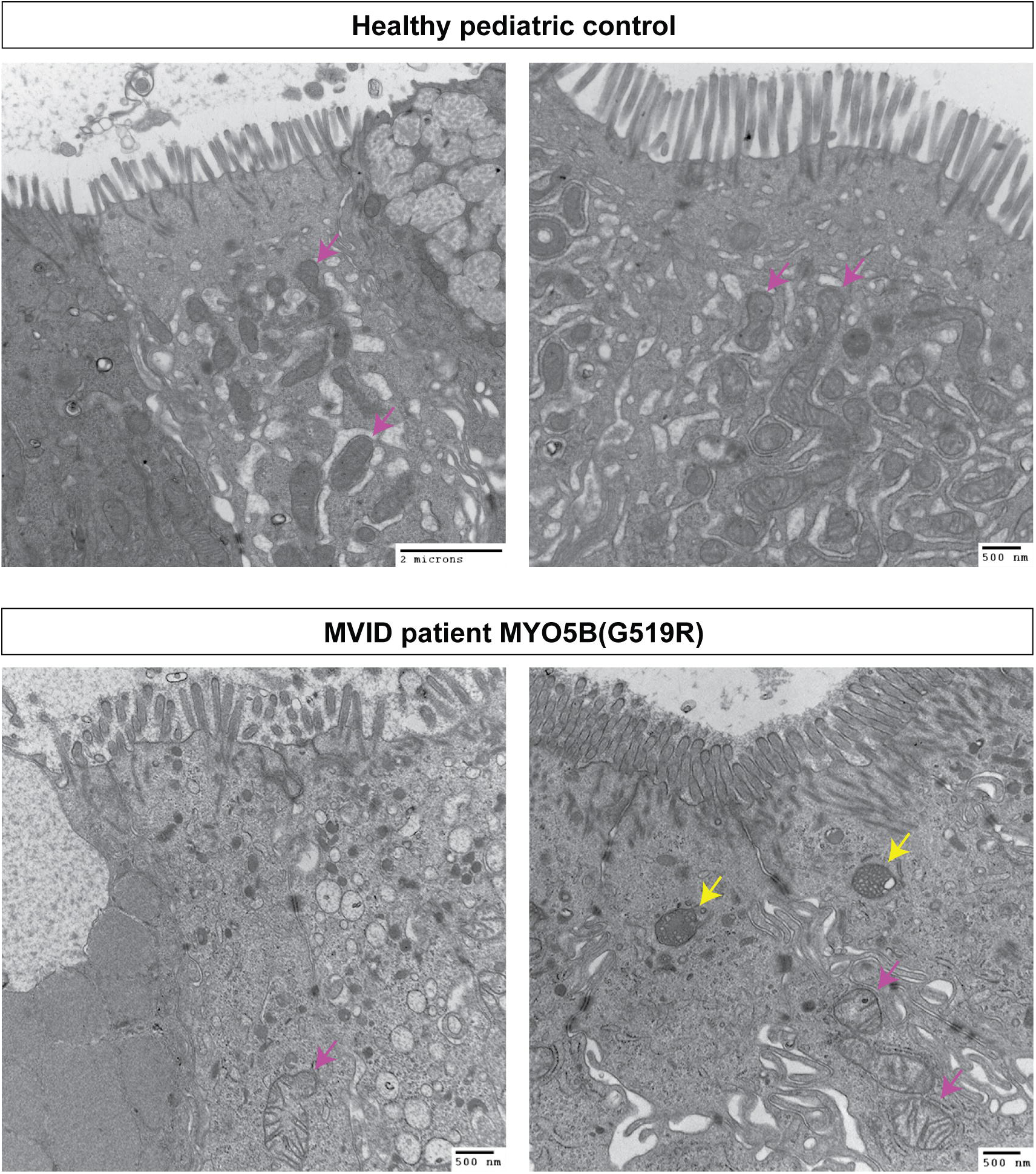
Disrupted mitochondria in MVID patient biopsy. Duodenal biopsies of healthy control and a MVID patient who possesses a compound heterozygous mutation in MYO5B (7). Control tissues harbor dense mitochondria including organized cristae (pink arrows). In the MVID patient tissues, disorganized microvillus structure of enterocytes is associated with abnormal formation of mitochondria (pink arrows) and multivesicular bodies (yellow arrows).

The mucosal morphological changes observed in MYO5B-deficient mice could be due to malnutrition after the loss of apical nutrient transporters. To compare these epithelial phenotypes in MYO5B-deficient mice to those in healthy mice under starvation conditions, control mice underwent food restriction for 48 hours. The fasted mice lost an average 23% of original body weight, but no signs of diarrhea or distress were observed. In the jejunum of fasted healthy mice, NDUFB8 staining showed a normal distribution in the cytoplasm, and the enterocytes possessed tall brush border structures (**Figure 3A** and **3B**). Thus, the abnormal distribution of mitochondria and immature brush border structures in MYO5BΔIEC enterocytes are likely caused by functional loss of MYO5B rather than following the nutrient malabsorption induced by transporter defects.

**Figure 3.**
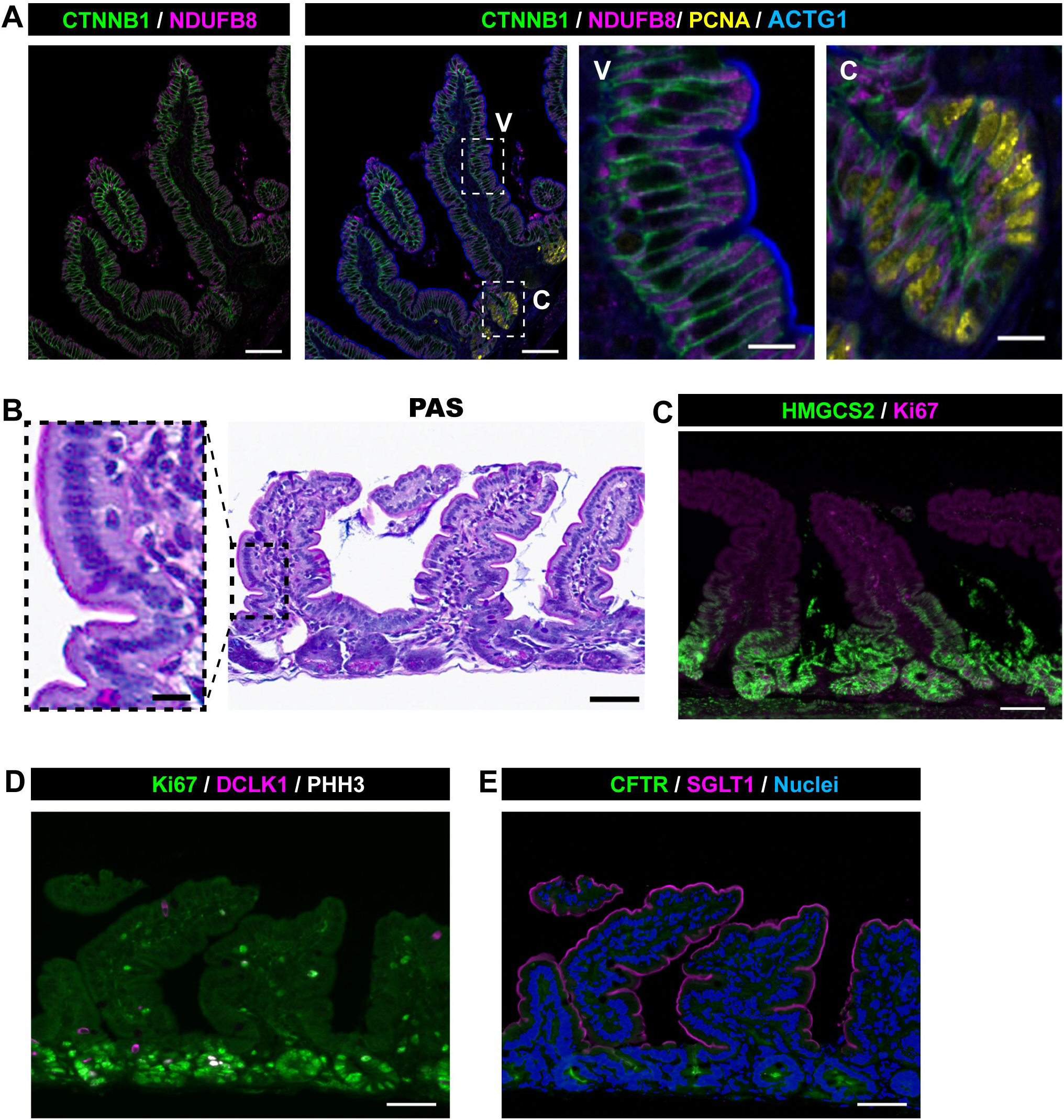
Jejunal tissue sections from fasted healthy mice. Four control mice were fasted for 48 hours, and their epithelial phenotypes are compared to those in MYO5B deficient mice. (A) Representative images of immunostaining for complex-I protein, NDUFB8. V=villus and C=crypt regions. (B) PAS staining shows established brush borders. (C) Immunostaining for a ketogenic enzyme, HMGCS2. (D and E) DCLK1+ tuft cell frequency and apical localization of SGLT1 and CFTR are similar to those of fed control tissues. Scale bars = 50 µm and 10 µm (insets).

### MYO5B loss alters cellular lipid metabolic pathways

The epithelial cell differentiation pathway closely interacts with epithelial cellular metabolism, in particular the fatty acid oxidation (FAO) pathway in stem cells (38–40). Utilizing our previous RNA-seq dataset of jejunal epithelial cells from MYO5BΔIEC mice (GSE139302) (12), we identified significant alterations in cellular metabolic pathways induced by MYO5B loss (**Figure 4A**). Transcription of genes that mediate the FAO, carnitine metabolism, and gluconeogenesis pathways were significantly downregulated. Conversely, MYO5B loss significantly upregulated genes that mediate fatty acid synthesis and lipogenesis (**Figure 4A**).

**Figure 4.**
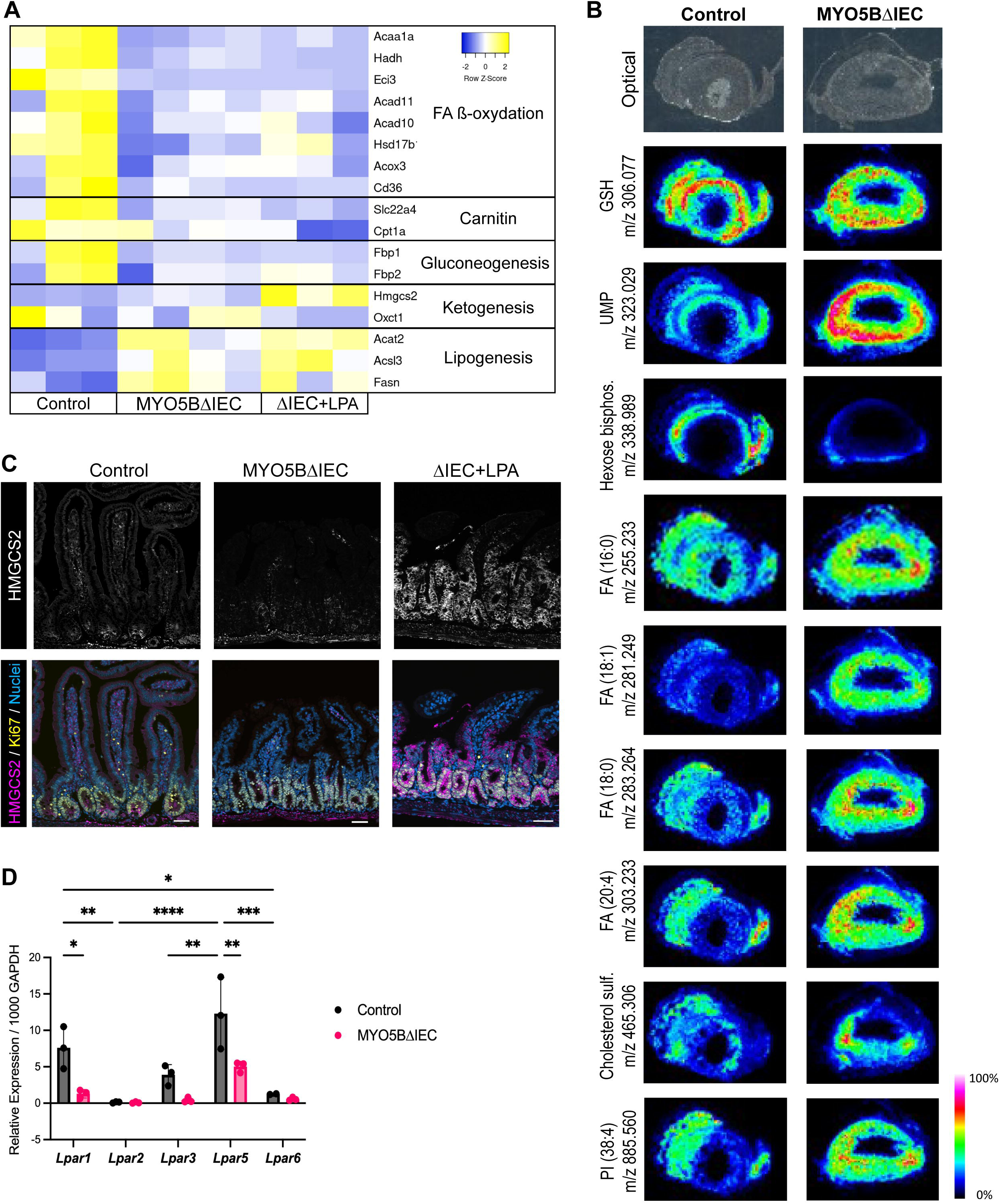
MYO5B loss and LPA treatment alters cellular energy metabolic pathways in mouse small intestine. (A) Transcriptional analysis in intestinal epithelial cells indicates significantly altered energy metabolic pathways in MYO5B deficient (MYO5BΔIEC) mice compared to control mice. LPA treatment on MYO5BΔIEC mice significantly increased the transcription of the rate-limiting ketogenic enzyme *Hmgcs2* compared to vehicle-treated group. (B) Imaging mass spectrometry was performed in fresh tissues of jejunum from 3 mice of each genotype, and representative images are shown as heatmaps. Optical images show morphology of horizontal sections of intestine with luminal space in the center. Derivatives of long-chain fatty acids and cholesterol are accumulated in MYO5BΔIEC mouse tissues, particularly in the villi. (C) Immunostaining for HMGCS2 and Ki67 in mouse jejunum. Daily LPA treatment increases the expression of HMGCS2 in the jejunal epithelial cells of MYO5BΔIEC mice. Scale bars = 50 µm. (D) Relative mRNA expression of LPA receptor (*Lpar*) subtypes in intestinal epithelial cells isolated from control and MYO5BΔIEC mice. Mean ± S.D. and each datapoint represents each mouse. \**P* < 0.05, by ANOVA with Tukey multiple comparisons. *n* = 3 mice per group.

To confirm the altered fatty acid metabolism induced by MYO5B loss, we employed imaging mass spectrometry (IMS) on mouse jejunal tissues. As shown in **Figure 4B**, the distribution of mass spectra of several metabolites was distinguishable between crypt and villus regions. Glutathione (GHS) was more abundant in crypt cells than in the villi of both control and MYO5B deficient tissues. An intermediate glycolysis metabolite, hexose bisphosphate, was less intense in MYO5B-deficient tissues than control, indicating decreased cellular metabolism in the MYO5B-deficient cells. Consistent with the RNA-seq results, the accumulation of derivatives of long-chain fatty acid and cholesterol were observed in MYO5B-deficient villi compared to control mouse tissues (**Figure 4B**). Uridine monophosphate (UMP) accumulation was identified in MYO5BΔIEC tissues, suggesting that MYO5B loss also influenced nucleotide metabolism.

Together, the activation of lipogenesis coupled with a decrease in FAO is induced by MYO5B inactivation in intestinal epithelial cells.

### LPA treatment upregulates ketogenesis pathway

We have previously demonstrated the therapeutic effect of lysophosphatidic acid (LPA) treatment on epithelial differentiation defects in MYO5B deficient mice (12, 13). Based on the transcriptional analysis, *in vivo* treatment with LPA(18:1) did not reverse the alterations in FAO or lipogenesis pathways observed with MYO5B loss (**Figure 4A**). Interestingly, LPA(18:1) treatment significantly upregulated *Hmgcs2*, a rate-limiting ketogenic enzyme, compared to vehicle-treated MYO5BΔIEC or healthy control (tamoxifen-untreated) mice (**Figure 4A**). To validate transcriptional changes in *Hmgcs2*, mouse tissues were immunostained for HMGCS2 (**Figure 4C**). The protein expression of HMGCS2 in healthy mouse jejunum is limited to the stem cells, as ketone bodies are essential for stem cell function (38). MYO5BΔIEC mouse jejunum showed expanded HMGCS2 expression in the proliferative cell zone, and systemic LPA treatment further increased the expression of HMGCS2 in both proliferating and differentiating epithelial cells (**Figure 4C**). The fasted mouse jejunum demonstrated a similar HMGCS2 expression pattern to the MYO5BΔIEC mice (**Figure 3C**). In addition to the stem cells, fasted jejunum also expressed HMGCS2 in proliferative cells and in enterocytes in the base of villi, where mature brush borders are established (**Figure 3B**). These observations suggest that HMGCS2 upregulation in proliferative epithelial cells of MYO5B-deficient intestine is a starvation phenotype, and that LPA treatment further increases ketone body production likely as an alternative energy fuel, resulting in the promotion of epithelial cell differentiation.

### Compound-1 specified the trophic effect of natural LPA through LPAR5 activation in two MVID model mice

To identify which LPA receptor (LPAR) signaling mediates the effect of LPA(18:1) treatment, *Lpar* mRNA expression was determined in jejunal epithelial cells of MYO5BΔIEC and control mice. Among LPAR1–6, LPAR4 was undetectable and all other LPAR subtypes were significantly lower in MYO5BΔIEC mice than control tissues (**Figure 4D**). However, LPAR5 expression was still prominent in MYO5B-deficient mouse intestine.

Based on the previous data with LPA(18:1) treatment and the presence of epithelial *Lpar5*, we next examined the effect of a synthetic agonist for LPAR5, Compound-1 (C_20_H_42_NaO_5_PS, mw: 448.57) (24). This chemical is highly water soluble (> 3.5 mg/ml) and has higher affinity to human LPAR5 than natural LPA(18:1) (24). Adult MYO5BΔIEC mice received intraperitoneal injection of Compound-1 (1 mg/kg) once a day following tamoxifen induction of MYO5B deletion (**Figure 5A**). The absence of MYO5B expression in epithelial cells was confirmed by immunostaining with or without Compound-1 treatment in MYO5BΔIEC mice (Supplementary Figure 1). To compare the treatment efficacy between MYO5B knockout and the MYO5B point mutation at G519R, which has been identified in an MVID patient (7), *Vil1-Cre^ERT2^;Myo5b^flox/G519R^* (MYO5B(G519R)) mice were treated with tamoxifen and Compound-1 in the same manner. Vehicle treated MYO5B(G519R) mice showed higher morbidity as 7 of 24 mice did not survive 4 days post tamoxifen induction, compared to MYO5BΔIEC mice (3 out of 31).

**Figure 5.**
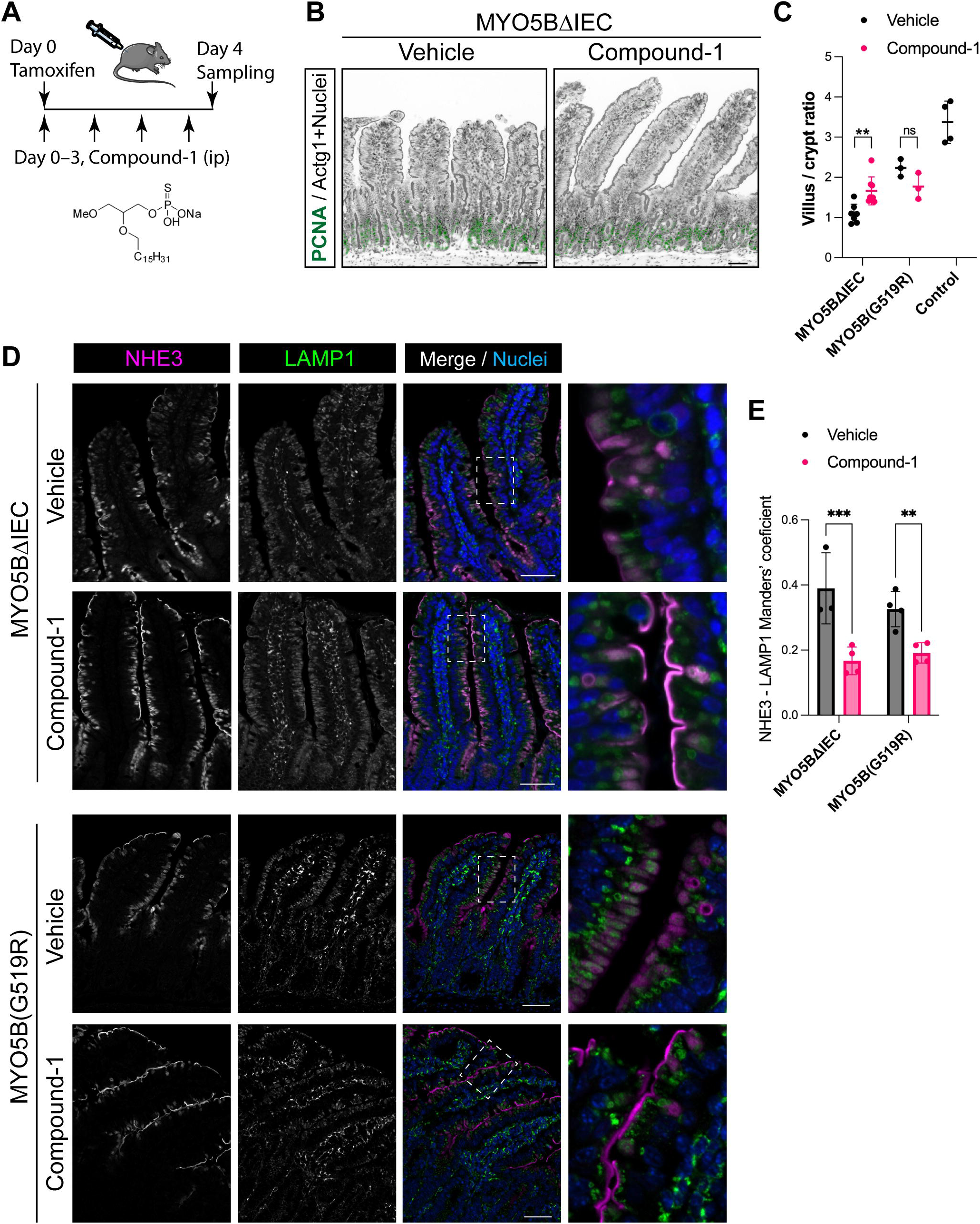
LPAR5 agonist, Compound-1, recapitulates the trophic effect of natural LPA. (A) Compound-1 structure and experimental design of mouse treatment. (B) Immunostaining for PCNA as a proliferative crypt marker, and ACTG1 for epithelial actin. (C) Villus/crypt ratio in jejunum of MYO5BΔIEC and MYO5B(G519R) mice. Bars indicate median values and each datapoint represents an average value of 10 regions in each mouse (*N* = 3–5 mice). (D) Immunostaining for NHE3 and LAMP1, a lysosomal marker, in the MYO5BΔIEC mouse jejunum. In vehicle-treated mice, NHE3 is diffuse in cytoplasm of epithelial cells or localized in lysosomes. Compound-1-treated mice demonstrate the separation of NHE3 from lysosomes and the re-establishment of the apical localization of NHE3. (E) Colocalization analysis of NHE3 and LAMP-1. Compound-1 significantly reduced the NHE3 accumulation in lysosomes. Mean ± S.D. and each datapoint represents the average of 5 regions in each mouse (*N* = 3–4 mice). Scale bar = 50 µm. \*\**P* < 0.01, \*\*\**P* < 0.001 by two-way ANOVA with LSD test.

The villus/crypt ratio in MYO5BΔIEC mouse jejunum was significantly improved by daily intraperitoneal treatment with Compound-1, indicating the increase in nutrient absorptive area (**Figure 5B, 5C and Figure 6A H&E**). Proliferative cells and mitotic cells of crypts were immunostained on jejunal sections for Ki67 and PHH3, respectively (**Figure 6A and 6B**). Compound-1 had no significant effect on either proliferative crypt length or PHH3+ cell numbers, suggesting that the improvement of villus/crypt ratio resulted from an increase in villus length. On the other hand, the villus/crypt ratio of MYO5B(G519R) mice did not show significant change with Compound-1 treatment (**Figure 5C, 6A and 6B**). These observations suggest that LPAR5 activates alternate cellular signaling pathways depending on MYO5B functional status, which is completely absent in MYO5BΔIEC mice, while the mutant MYO5B(G519R) protein may influence epithelial cell functions.

**Figure 6.**
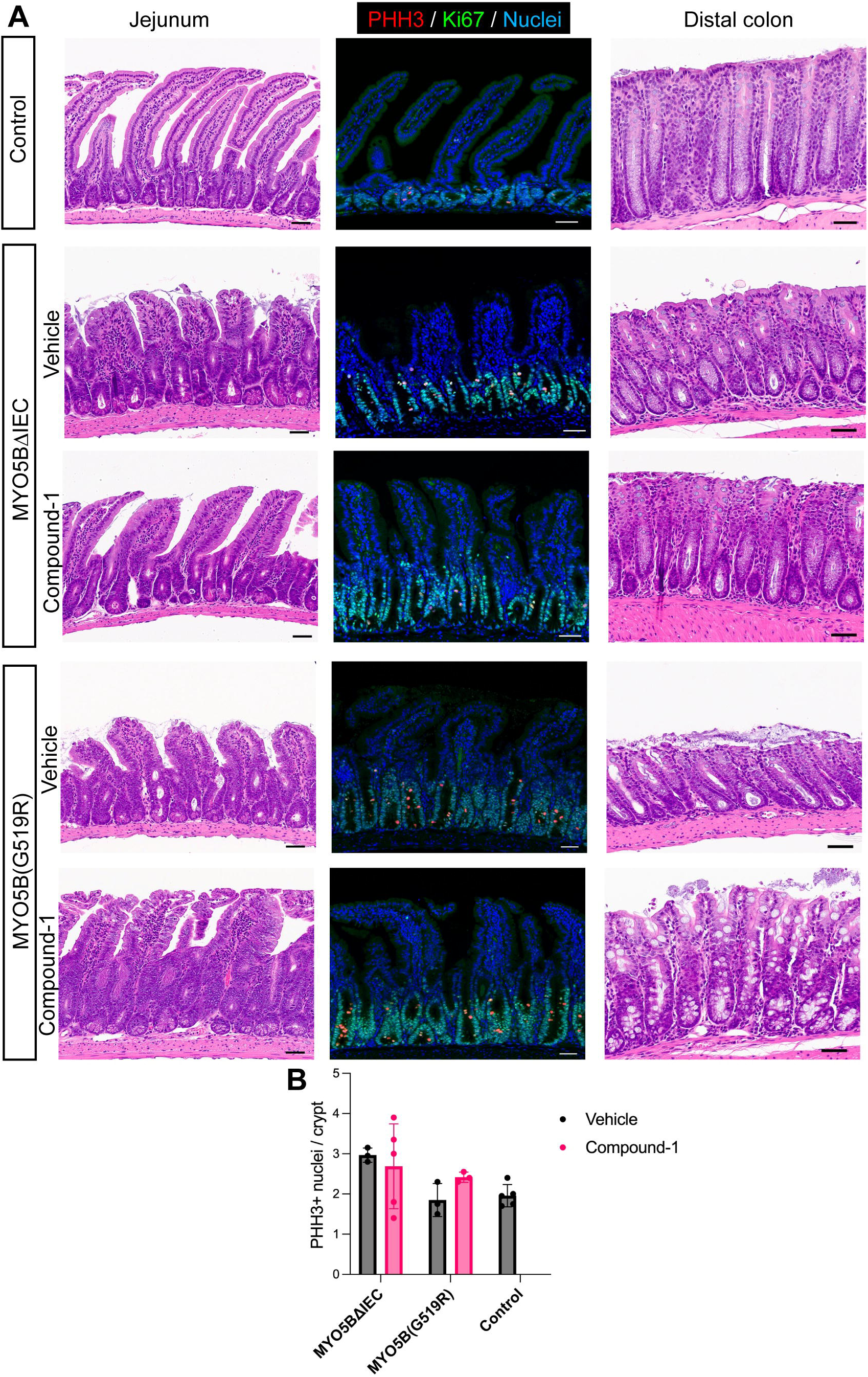
Histology and proliferative cell population in crypts. (A) H&E-stained jejunal and colonic tissue sections and immunostaining for proliferation markers. MVID model mice demonstrate expanded crypts and blunted villi in the jejunum and the dilation of colonic crypts. Compound-1 treatment improves villus morphology of MYO5BΔIEC mice, but does not remarkably alter MYO5B(G519R) villi. Mitotic marker, PHH3, and a proliferating marker, Ki67, in jejunal tissues demonstrate elongated crypt area in both MVID models. **(B) Quantification of mitotic cells in the jejunum.** PHH3+ nuclei per crypt were counted in 20 regions of each mouse section. Each datapoint on the graph indicates the averaged value of each mouse. *N* = 3–5 mice per group. No significant difference was detected by ANOVA. Scale bars = 50 µm.

Na^+^/H^+^ exchanger (NHE)3 is an important brush border protein for sodium absorption. The immunoreactivities of NHE3 were mislocalized from the brush border and mostly localized with a lysosomal marker, LAMP1, in the subapical space of vehicle-treated MYO5BΔIEC and MYO5B(G519R) enterocytes, indicating abnormal protein degradation of these mis-trafficked transporters (**Figure 5D**). Compound-1 treatment improved NHE3 localization to the brush border and significantly reduced the colocalization of NHE3 and LAMP1 by approximately 50% (**Figure 5E**), indicating that LPAR5 activation ameliorates brush border maturation in both MVID mouse models: MYO5B deficient mice and the mice with a point mutation at MYO5B(G519R).

Similarly, Na^+^-dependent glucose transporter (SGLT)1 is a crucial apical protein for water absorption and is mislocalized in MVID intestinal tissues. Small intestine of both MYO5BΔIEC and MYO5B(G519R) mice demonstrated cytoplasmic concentrations of SGLT1, colocalized with LAMP1 (**Figure 7A**). SGLT1 immunostaining in Compound-1-treated mouse tissues were significantly separated from lysosomes and localized on brush border (**Figure 7A and 7B**). SGLT1 expression is limited to mature enterocytes of villi more specifically than NHE3. To quantify cellular SGLT1 distribution in the enterocytes, we developed a cell segmentation analysis script. As shown in **Figure 7C** Overlay, individual epithelial cells were segmented utilizing general membrane markers, gamma-actin (ACTG1) and beta-catenin (CTNNB1). Next, localization of SGLT1 staining was defined in apical membrane versus cytoplasm areas of each segment, and the intensity ratio of the membrane to cytoplasm was visualized as a heatmap (**Figure 7C**). In each Swiss roll, more than 30,000 epithelial cells were analyzed and the mean values of each treatment group (*N* = 3–4 mice per group) were shown as histogram in Figure 4D. Mean values of SGLT1 ratio were shifted toward the membrane (larger values) in Compound-1-treated MYO5BΔIEC and MYO5B(G519R) mice compared to vehicle-treated groups (**Figure 7D**).

**Figure 7.**
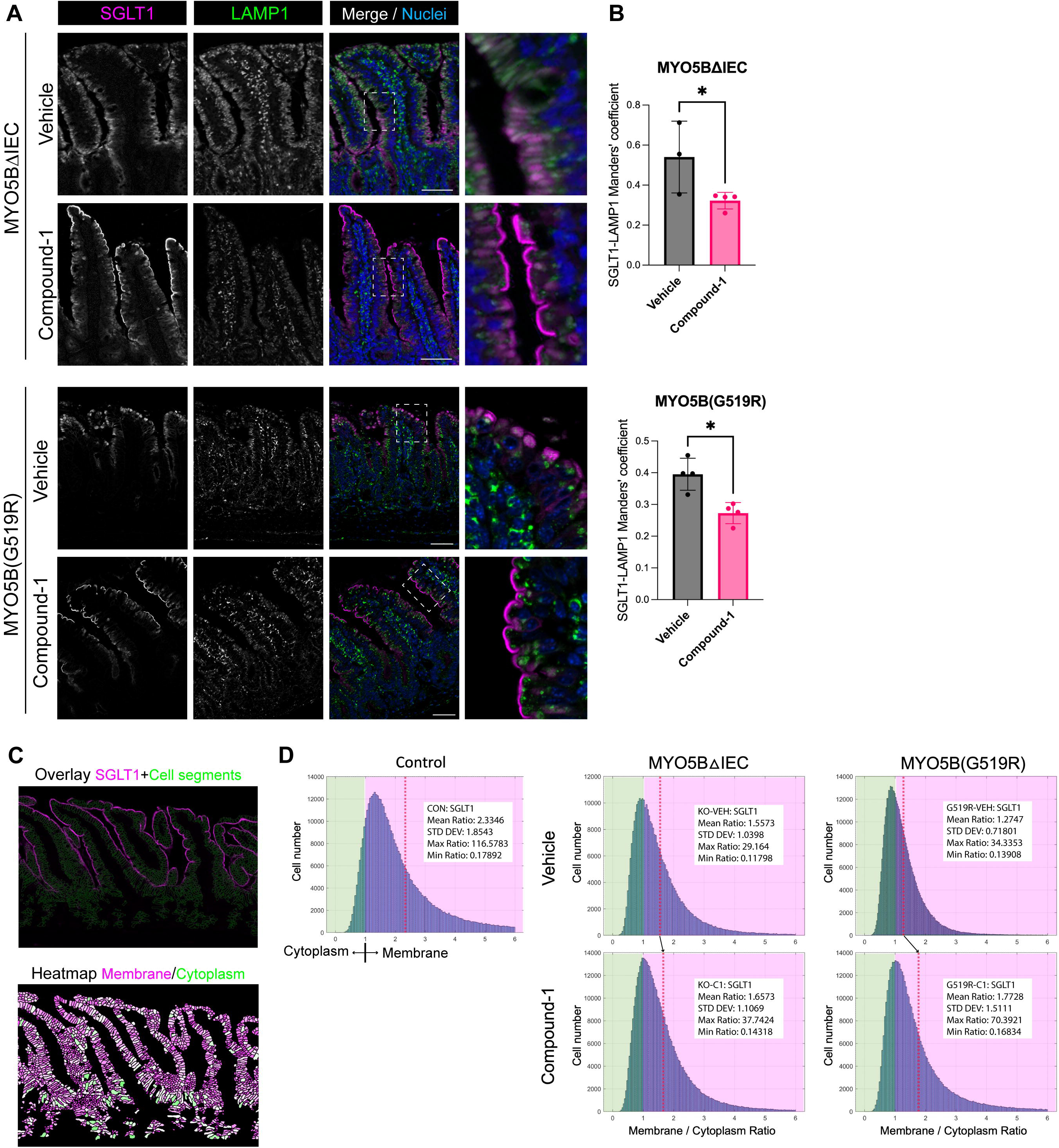
Membrane SGLT1 localization is enhanced by Compound-1 treatment in MVID model mice. (A) Immunostaining for SGLT1 and LAMP1 in jejunum of MYO5BΔIEC and MYO5B(G519R) mice. (B) Colocalization analysis of SGLT1 and LAMP-1. \**P* < 0.05 by Mann-Whitney test. (C) Representative images of cell segmentation analysis in control tissue sections. (D) Histograms representing SGLT1 intensity ratios of all identified epithelial cells from jejunum of four mice in each group. Red dash lines indicate the mean values. Compound-1 treatment shifts the mean ratio in MVID model mice towards membrane localization.

Immunostaining pattern of the mitochondrial complex-I marker, NDUFB8, was investigated in Compound-1 treated MYO5BΔIEC and MYO5B(G519R) mice. Consistent with a partial improvement of NHE3 and SGLT1 presentation in the brush border after Compound-1 treatment, mitochondrial distribution in subapical area was partially rescued both in MYO5BΔIEC and MYO5B(G519R) mouse intestine (**Figure 8**). Brush border structure was visualized by co-immunostaining for villin. The improved brush border in the treated mice showed long, dense microvilli and NDUFB8 signals in the cellular space adjacent to the microvilli (**Figure 8** white squares). The Compound-1-treated mouse tissues included unhealthy enterocytes with thin brush border, and these cells demonstrated diminished mitochondrial signal in subapical area (**Figure 8**, yellow squares). These observations indicate the correlation of brush border status and mitochondrial activity in the intestinal epithelial cells.

**Figure 8.**
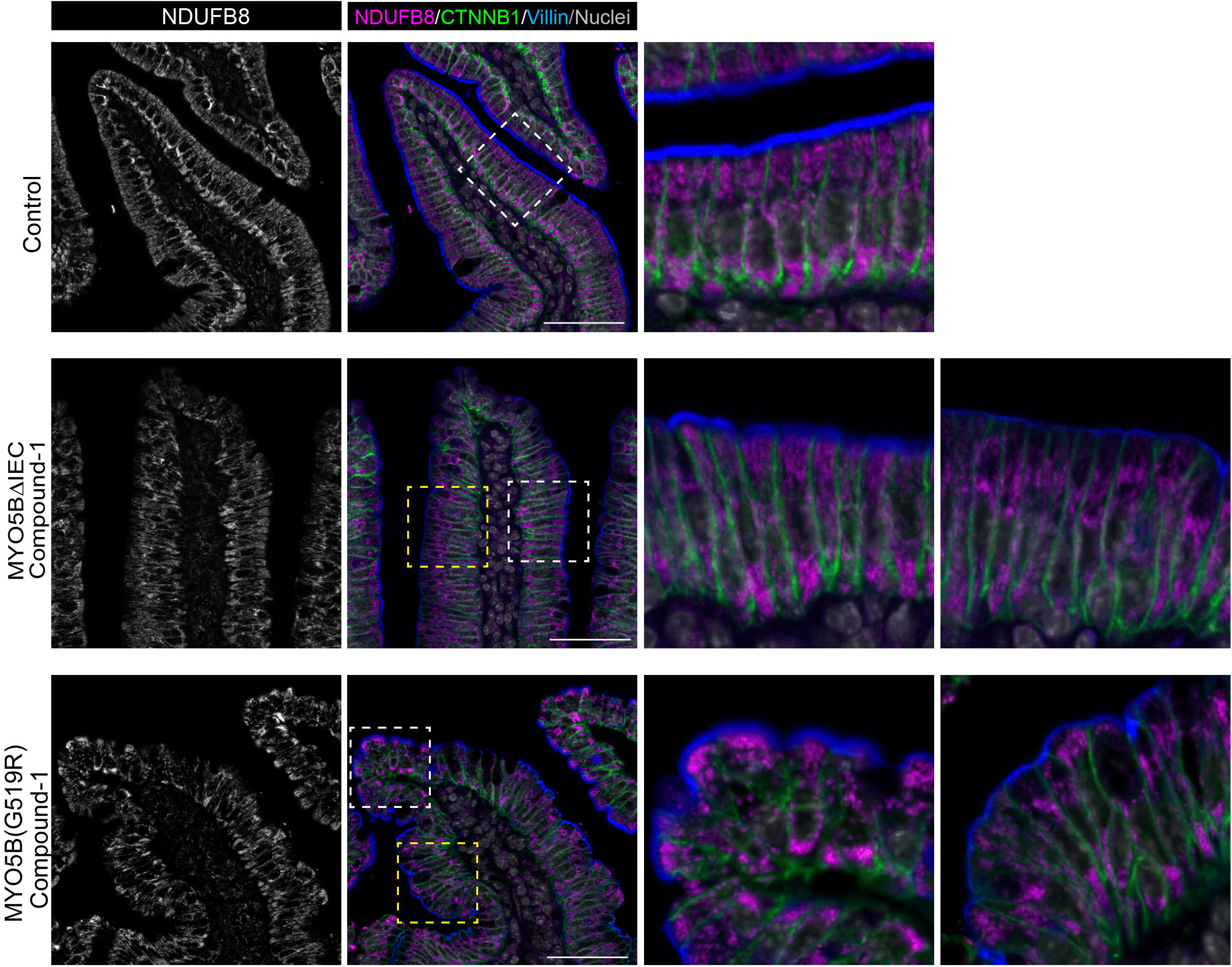
Mitochondrial distribution is partially rescued by Compound-1 treatment in MVID model mice. Jejunal sections of MYO5BΔIEC and MYO5B(G519R) mice were immunostained for NDUFB8 and villin after daily Compound-1 treatment. Control tissues show uniformed dense brush border in villi, and abundant subapical mitochondria close to the brush border. Both MVID model mice demonstrate some improved enterocytes that possess thick brush border and intense mitochondrial signal in the subapical area (white squares). However, the other part of villi had unhealthy enterocytes that show thin brush border and diminished mitochondrial signal in the subapical area (yellow squares). Scale bars = 50 µm.

Despite the improvements of enterocyte transporter presentation on the brush border, body weight loss was observed following tamoxifen injection in both MVID model mice (**Figure 9A**). Only male mice with MYO5B(G519R) mutation showed a significant improvement of body weight by Compound-1 treatment. To verify the immunohistological outcome of transporters, sodium absorptive function was assessed in jejunum by using Ussing chamber system, SGLT1-mediated *I*_sc_ was significantly increased by Compound-1 treatment both in male (triangle datapoints) and female (circle datapoints) mice of both MYO5BΔIEC and MYO5B(G519R) strains compared to vehicle-treated mice (**Figure 9B**). However, SGLT1 activity in the treated mouse tissues were still significantly lower than that in healthy control littermates. Electrogenic chloride secretion, which induces water secretion into the lumen, was also assessed. Calcium- and cAMP-induced secretion were analyzed by applying carbachol (CCh) and forskolin (FK), respectively. Secretory response to cAMP, but not calcium, was significantly increased in vehicle-treated MYO5B(G519R) mice than controls (**Figure 9C, 9D**). CFTR-dependent portion of the secretory response was further assessed by applying a CFTR inhibitor, (R)-BPO-27 (**Figure 9E**). MYO5BΔIEC tissues showed significantly higher CFTR activity than controls, consistent with our previous observations. Compound-1 treatment had no effect on secretory responses or CFTR activity in either MVID model mice, suggesting that LPAR5 activation does not affect secretory symptoms in MVID.

**Figure 9.**
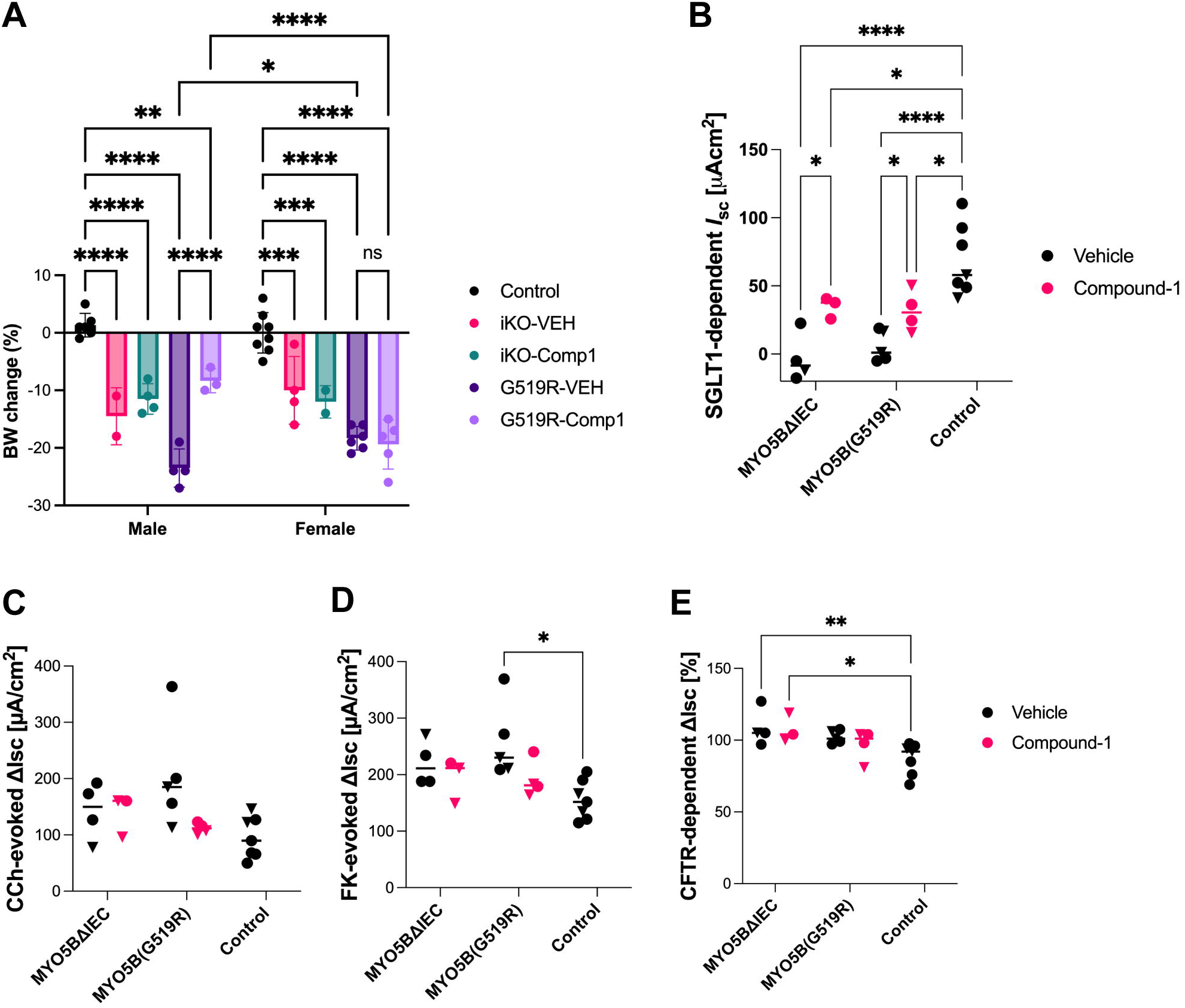
Differential effect of Compound-1 treatment on body weight loss and epithelial transporter function. (A) Change in body weight on day 4 post-tamoxifen compared to the original weight on day 0. Each datapoint indicates the value of each mouse. Among genotype and sex groups, only MYO5B(G519R) male mice show a significant improvement of body weight by Compound-1 treatment. \**P* < 0.05, \*\**P* < 0.01, \*\*\**P* < 0.001 by two-way ANOVA with Tukey’s test. **(B) SGLT1 activity in Ussing chambered jejunum.** Change in short-circuit current (*I*_sc_) was measured in response to a SGLT1 inhibitor, phlorizin. Compound-1 treatment significantly increased SGLT1 function both in MYO5BΔIEC and MYO5B(G519R) mice. Circle datapoints indicate female mouse tissues and triangle datapoints indicate male mouse tissues. \**P* < 0.05, \*\*\*\**P* < 0.0001 by two-way ANOVA with Tukey’s test. **(C-E) Secretory responses in the jejunum.** Chloride and water secretion was sequentially stimulated with carbachol (CCh; C) and forskolin (FK; D). CFTR dependency in secretory state was measured by a CFTR inhibitor (R)-BPO (E). There is no statistical difference between vehicle vs. Compound-1 treatment in each genotype. \**P* < 0.05, \*\**P* < 0.01 by two-way ANOVA with Tukey’s test.

Membrane-bound mucins, such as MUC13, are important for intestinal barrier function (41). We previously reported that the deficiency of Rab11-FIP1 impairs MUC13 localization in the colon, correlating to an increased susceptibility to experimental mucosal inflammation in mice (42). As a mature brush border marker, MUC13 localization was investigated in the present Compound-1 treated mice. MUC13 is localized to the tips of microvilli in control small intestine and colon, however, both MYO5BΔIEC and MYO5B(G519R) mouse tissues demonstrated strong MUC13 signals in the cytoplasm of epithelial cells throughout the small intestine and colon (**Figure 10**). In Compound-1-treated MYO5BΔIEC and MYO5B(G519R) mouse tissues, most MUC13 signals were still localized to the cytoplasm despite the improvement of villus morphologies (**Figure 10**). These observations suggest that the apical sodium transporters and MUC13 trafficking pathways are functionally MYO5B-dependent, however, LPAR5 activation alone was not sufficient to re-establish MUC13 localization to the microvilli tips.

**Figure 10.**
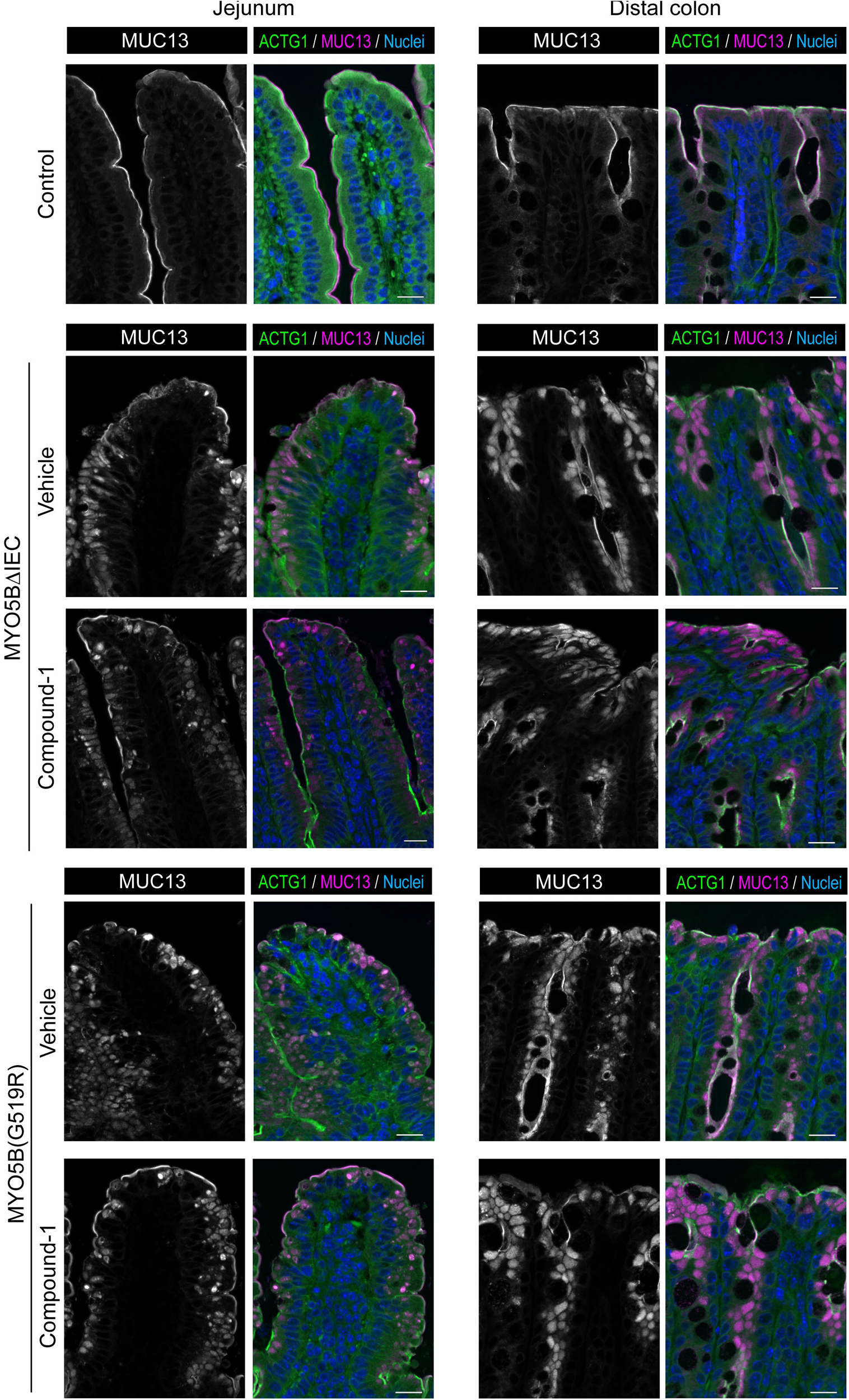
Lack of effect of Compound-1 on MUC13 localization in MVID model mice. Immunostaining for MUC13 and ACTG1 in jejunal tissues of control, MYO5BΔIEC, and MYO5B(G519R) mice treated with Compound-1 or vehicle. In control jejunum and distal colon, MUC13 (red) is localized to the apical tips of microvilli above actin filaments (green). Both MYO5B loss-of-function mouse tissues demonstrate strong MUC13 immunostaining in cytoplasm of jejunal and colonic epithelial cells. The mislocalization of MUC13 is not remarkably altered by Compound-1 treatments in MYO5BΔIEC or MYO5B(G519R) mice. Scale bars = 20 µm.

### LPAR5 activation by Compound-1 ameliorates tuft cell differentiation in MVID models

We recently reported that MYO5B loss disrupts the epithelial cell lineage differentiation through an imbalance of epithelial Wnt/Notch signaling (13). To assess epithelial cell differentiation competence, the population of a sensory epithelial cell, the tuft cell, was quantified in MYO5BΔIEC mice with different LPAR agonist treatments. Similar to natural LPA(18:1), Compound-1 enhanced tuft cell differentiation in the MYO5BΔIEC mouse intestine (**Figure 11A**). Other synthetic LPAR agonists had no significant effect, including UCM-05194 (10 mg/kg) or GRI977143 (3 mg/kg), selective agonists for LPAR1 or LPAR2, respectively. Tuft cell numbers in MYO5B(G519R) mouse intestine varied more widely than that in MYO5BΔIEC mice and were not increased by Compound-1 treatment (**Figure 11B**).

**Figure 11.**
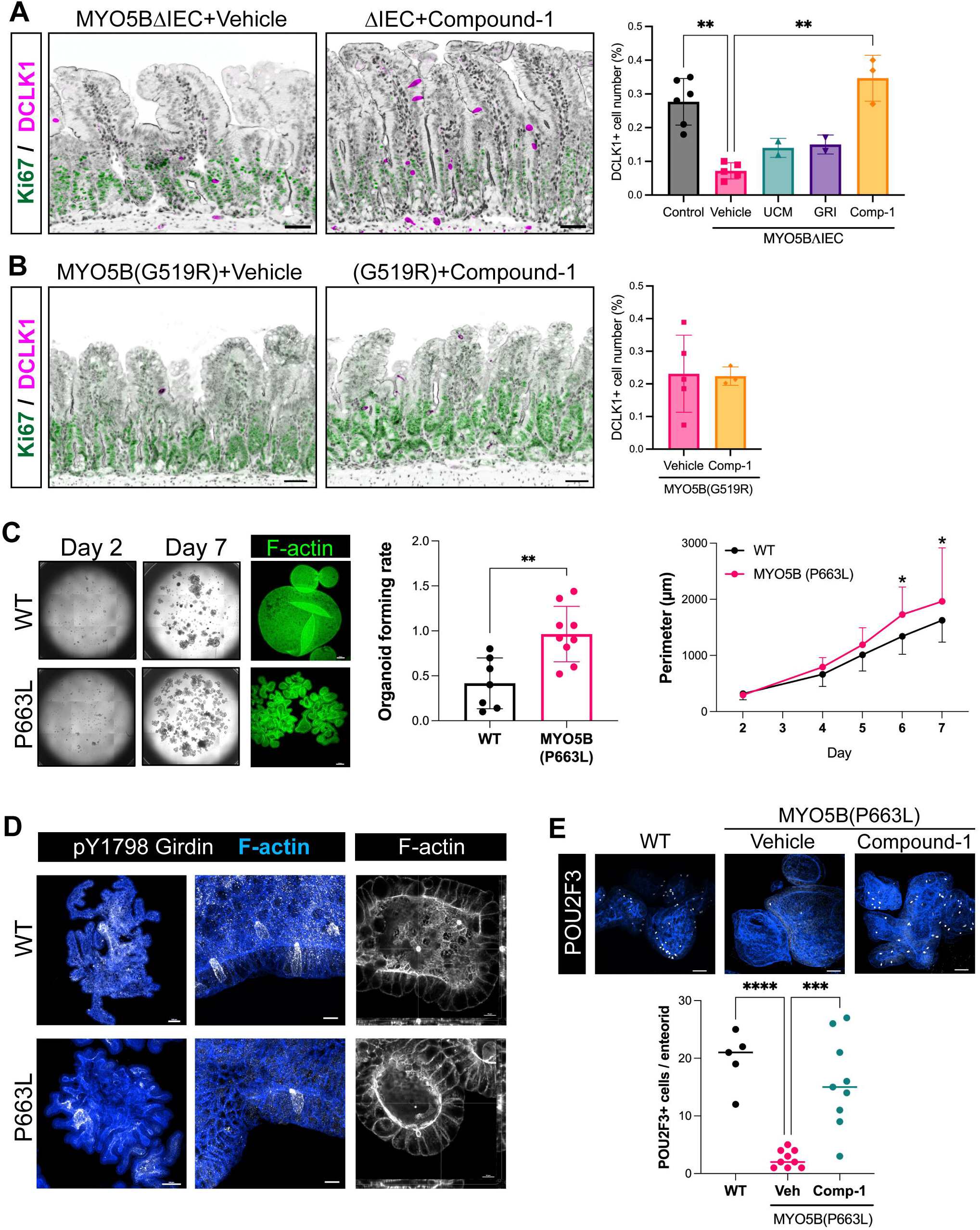
LPAR5 activation ameliorates tuft cell differentiation in MVID models. (A and B) Immunostaining for DCLK1 in the jejunum of MYO5BΔIEC (A) and MYO5B(G519R) (B) mice treated with Compound-1 or vehicle. DCLK1+ tuft cells (magenta) do not express the proliferative marker, Ki67 (green). Counterstaining for ACTG1 and nuclei is shown in black as an inverted color. Scale bars = 50 µm. Tuft cell number per mucosal cell number is determined in whole jejunal Swiss rolls by digital image analysis. Each datapoint indicates a value from each mouse. \*\**P* < 0.01 by Kruskal-Wallis test with Dunn’s multiple comparisons. (C) Enteroids were generated from jejunal crypts of wild-type (WT) and genetically engineered pig with the point mutation MYO5B(P663L) and expanded in Human Organoid Growth Medium. Whole well images were analyzed to determine organoid forming efficacy and perimeters in wild type (WT) and MVID models. Scale = 100 µm. \**P* < 0.05 by two-way ANOVA. (D) Whole-mount immunostaining for tuft cell markers, pY1798-Girdin and POU2F3, and phalloidin staining for F-actin. (E) Differentiation medium was supplemented with Compound-1 (100nM) or vehicle for 5 days. MYO5B(P663L) enteroids have less tuft cells compared to WT, and Compound-1 treatment significantly increased tuft cell differentiation. Each datapoint indicates each enteroid. \*\*\**P* < 0.001 by ANOVA with Dunnett’s multiple comparisons.

The small intestine from fasted control mice demonstrated normal localization of SGLT1 and CFTR on the brush borders (**Figure 3D**). DCLK1+ mature tuft cells were distributed among villi and crypts (**Figure 3E**), similar to previously reported patterns (43). These observations support the concept that internalized brush border transporters and reduced tuft cell differentiation were specifically induced by MYO5B loss in progenitor cells.

The promotion of tuft cell differentiation by Compound-1 was tested in pig enteroid cultures that possess an MVID patient-modeling point mutation at MYO5B(P663L), which is the orthologue of human P660L (10). The MVID model enteroids generated from the jejunum of this mutant model demonstrated a higher enteroid forming rate than wild-type (WT) jejunal enteroids (**Figure 11C**). Pig enteroid tuft cells were differentiated in IntestiCult™ organoid differentiation medium (ODM) for 6 days and immunostained for a tuft cell marker, phosphorylated Girdin (pY-1798) (32) and the key transcription factor for sensory cell lineages, POU2F3 (44). DCLK1, which is widely used for intestinal tuft cell marker in rodent tissues did not stain pig or human tuft cells in enteroids. Immunostaining for pY1798-girdin in pig enteroids was identified in apical tuft structure and cytoplasm of tuft cells, which possess dense F-actin in microvilli (**Figure 11D**), similarly to that in human intestinal tuft cells (45). F-actin staining with phalloidin visualized inclusion formation in MYO5B(P663L) enteroids, indicating the differentiation of MVID-affected enterocytes (**Figure 11D**). Compared to WT pig enteroids, MYO5B(P663L) enteroids had significantly fewer POU2F3+ cells in each enteroid (**Figure 11E**). These results suggest that MYO5B loss-induced differentiation defects are not species-specific. Daily supplementation of Compound-1 (100 nM) into the ODM significantly increased tuft cell numbers in MYO5B(P663L) enteroids, suggesting that LPAR5 activation has cell-autonomous effects on MVID model enteroids (**Figure 11E**).

### Transcription signatures in MYO5B inactivated enteroids show the defects of cellular energy metabolism

To assess the epithelial cell-autonomous effects of MYO5B inactivation, bulk RNA-sequencing analysis was performed in enteroids that were generated from MYO5BΔIEC and MYO5B(G519R) mouse jejunum (GSE260706). The overall alteration pattern of gene transcripts showed high similarity between MYO5BΔIEC and MYO5B(G519R) enteroids (**Figure 12A** and **Supplementary Tables 1-3**), which is consistent with our pathological observations of *in vivo* mouse tissues (7).

**Figure 12.**
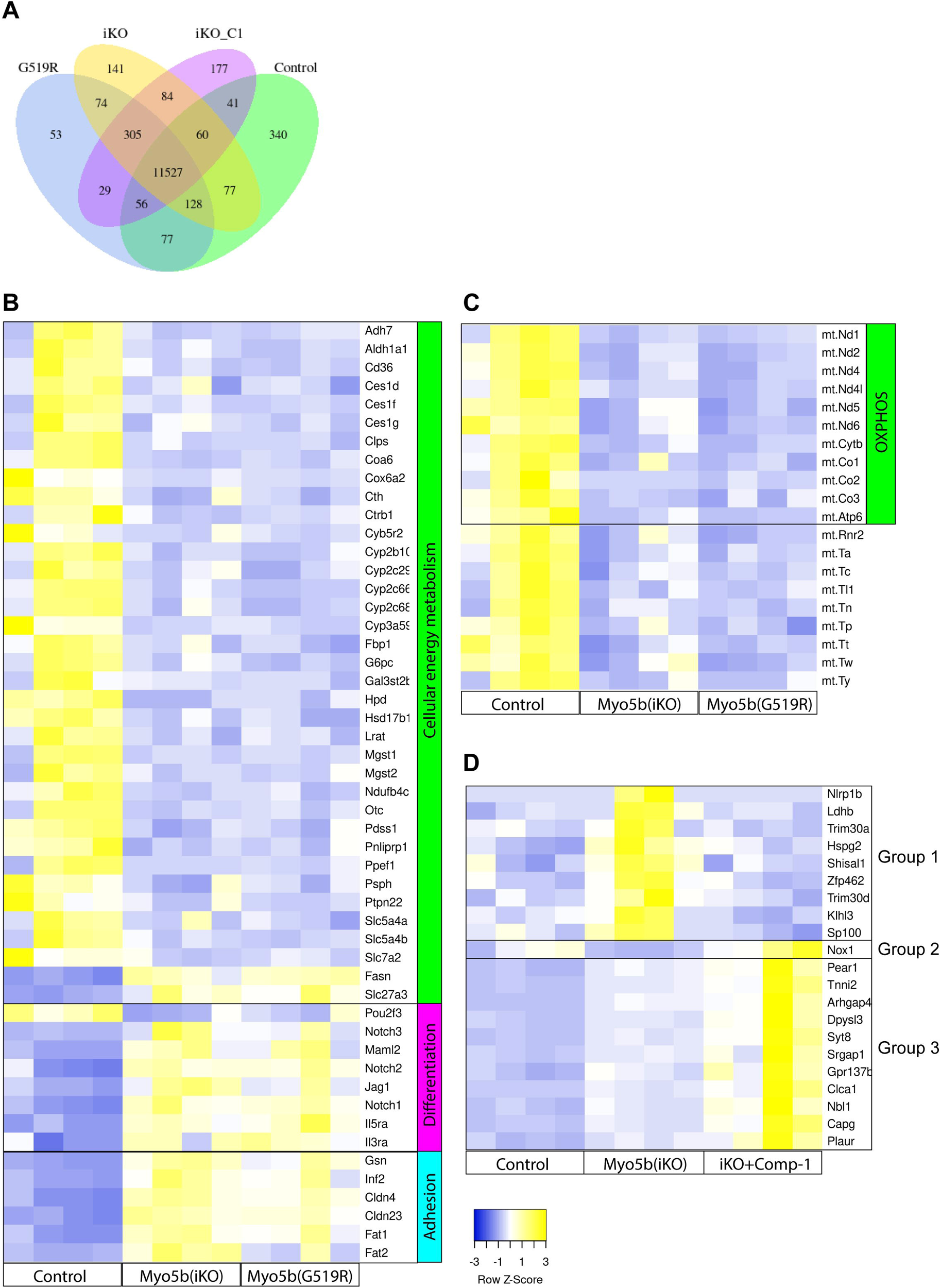
Transcription signatures in MYO5B inactivated enteroids show the defects of cellular energy metabolism. (A) Summary of differential expression genes in mouse enteroids. Four individual cultures from four mice of each group are analyzed; iKO represents MYO5BΔIEC, iKO_C1 is Compound-1-treated MYO5BΔIEC, G519R is MYO5B(G519R), and Control is healthy littermates. (B) Selected transcription signatures in enteroids generated from MYO5BΔIEC (iKO) and MYO5B(G519R) mice. (C) Mitochondrial gene transcriptions including most OXPHOS metabolic genes were significantly decreased in both iKO and MYO5B(G519R) enteroids. (D) MYO5B deficient enteroids were treated with Compound-1 (100 nM) or vehicle for 2 days. Group 1 genes are significantly increased by MYO5B loss and decreased by Compound-1 treatment. Only *Nox1* is decreased by MYO5B loss and reversed by Compound-1, while Group 3 genes are increased by MYO5B loss and further increased by Compound-1. *N* = 4 mice per each genotype.

MYO5BΔIEC enteroids were treated with Compound-1 (100 nM) for 2 days during differentiation to evaluate direct LPAR5 targets in enterocytes. *Lpar2*, *Lpar5*, and *Lpar6* transcription was detected in all samples and their transcripts were expressed at comparable levels among control, untreated and Compound-1-treated MYO5BΔIEC enteroids. Even though MYO5BΔIEC and MYO5B(G519R) enteroids were cultured in nutrient-rich medium and grown in a similar manner to control enteroids, numerous genes of energy metabolic enzymes were significantly downregulated by functional loss of MYO5B (**Figure 12B and 12C**). On the other hand, fatty acid synthesis genes, *Fasn* and *Slc27a3,* were significantly upregulated in both enteroid models similar to the transcriptional signatures in the epithelial cells from MYO5BΔIEC tissues (**Figure 12B**). Furthermore, 20 mitochondrial genes, including oxidative phosphorylation (OXPHOS) pathway, were significantly decreased in both MVID model enteroids, indicating that functional MYO5B loss directly impairs mitochondrial activity and cellular energy metabolism of the epithelial cells (**Figure 12C**). In contrast, *Hmgcs2* expression was comparable between control and MYO5B-deficient enteroids with and without Compound-1, suggesting that the ketogenic pathway is maintained under enteroid culture conditions unlike what is observed in mouse tissues.

Both MYO5BΔIEC and MYO5B(G519R) enteroids showed significantly downregulated *Pou2f3* expression and upregulated *Notch1*, *Notch2*, *Notch3*, *Jag1*, and *Maml2*, indicating that tuft cell lineage differentiation was likely blocked through abnormally enhanced Notch signaling. Other remarkably upregulated genes associated with functional MYO5B loss are implicated in a proinflammatory response and abnormal cell-cell adhesion, including *Mcam, Plaur, Fat1, Fat2, Inf2, Col4a, Col4a2*, *Cldn4*, and *Cldn23*, which were very low or at undetectable levels in control enteroids (**Figure 12B and Figure 13A**). Differentially expressed gene lists of enteroids were compared to the previous datasets of isolated epithelial cells from MYO5BΔIEC mouse jejunum (GSE139302) (12) (**Supplementary Table 4**). Mutual 661 genes were significantly up- or down-regulated by MYO5B loss in tissues and enteroid cultures (**Figure 13A, 13B, and Supplementary Table 5**). Those 661 genes included Notch signaling molecules and proinflammatory markers as listed above, suggesting that these cell-stress signaling pathways are dependent on cell-autonomous MYO5B function and independent from nutrient availability. Important chemokines, *Ccl9* and *Ccl25*, were markedly downregulated in both MYO5B deficient enterocytes and epithelial tissues, implicating the direct influence of MYO5B loss on epithelial-immune crosstalk through chemokine signaling. *In vitro* Compound-1 treatment significantly reversed several inflammation-related genes, such as *Nlrp1b*, *Ldhb*, *Hspg2*, and *Trim30*, suggesting that Compound-1 directly improves some epithelial cell deficits induced by MYO5B loss (**Figure 12D**).

**Figure 13.**
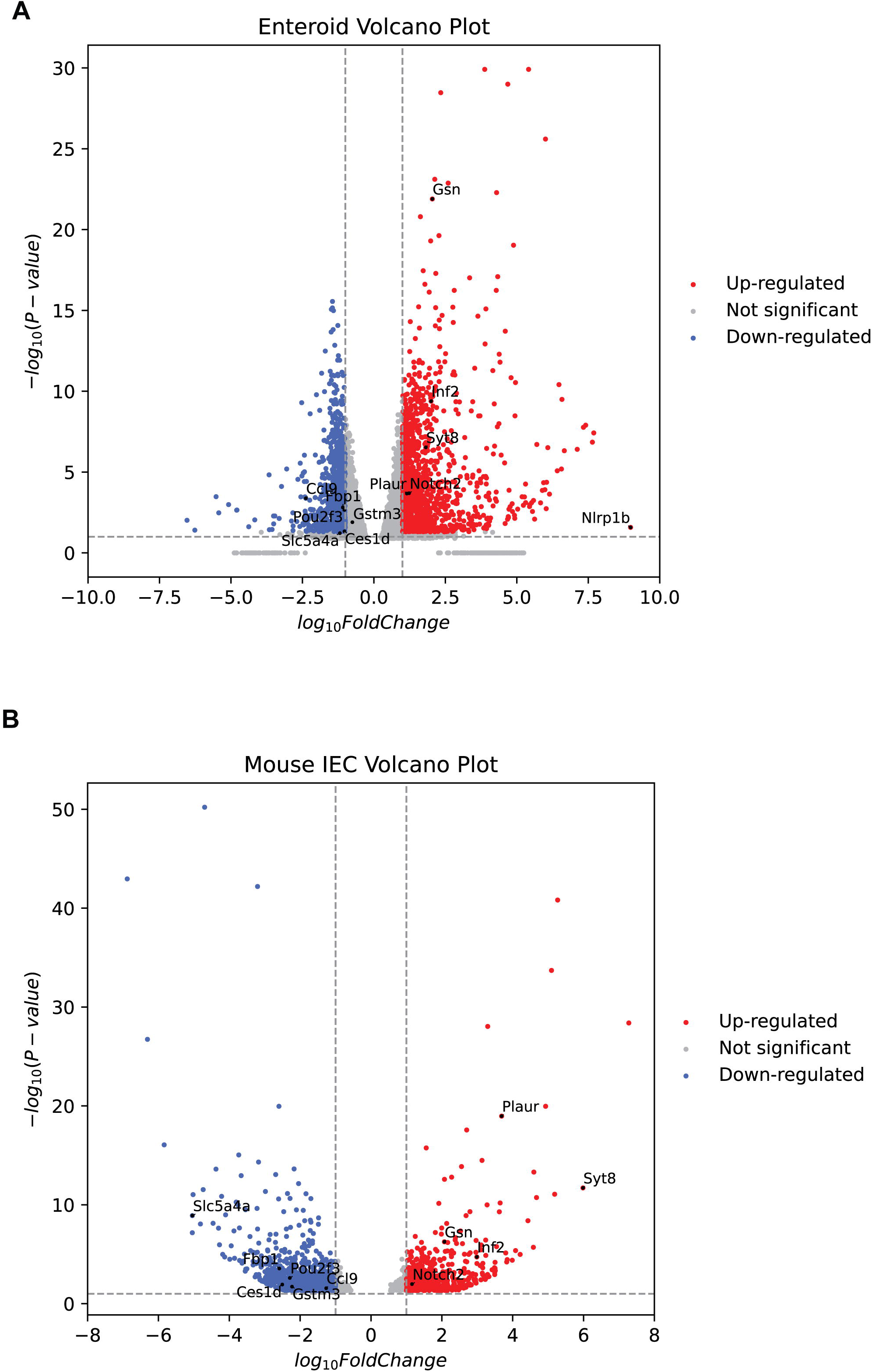
Differential expression of genes in MYO5BΔIEC enteroids (A) and epithelial cells isolated from jejunal tissues (B) compared to each control sample. Mutual genes of interest that are up- or down-regulated in MYO5B-deficient epithelial tissues and enteroids were marked as black dots.

## Discussion

### Impact of MYO5B loss on epithelial cellular metabolism

Augmenting absorptive function is a critical goal for the treatment of chronic diarrhea and malabsorption. However, enterocyte cellular development pathways have not been well considered as therapeutic targets of congenital diarrheal diseases. We have recently shown that the disrupted epithelial cell differentiation and enterocyte maturation in MYO5B-deficient intestine is likely another pathology that underlies the malabsorption symptoms of MVID (12, 13), in addition to MYO5B-mediated transporter trafficking defects investigated in differentiated cells (46, 47). The present study further indicates that fatty acid oxidation (FAO) and mitochondrial function of intestinal epithelial cells are significantly impaired by MYO5B defects in both mouse tissues and enteroid models. These metabolic defects are consistent between knockout and a point mutant (G519R) of MYO5B, which recapitulate MVID patients. Recent studies have revealed that FAO and mitochondrial metabolism are essential for stem cell maintenance and differentiation (35, 39), suggesting that MYO5B-mediated vesicle trafficking is involved in cellular metabolic pathways of intestinal stem cells.

The specific functions of MYO5B in intestinal progenitor cells have not been well characterized. Our TEM images demonstrated damaged mitochondria in crypt progenitor cells in mouse and patient tissues (**Figure 1A and 2**). Previous studies demonstrate that myosin V sequesters BCL-2-modifying factor (BMF) protein, which binds and inactivates mitochondria that leading to apoptosis in intestinal epithelial cells (48, 49). These studies suggest that MYO5B loss can indirectly impair mitochondria. Furthermore, in enteroid models of two MYO5B loss-of-function mice (MYO5B(G519R) and MYO5BΔIEC), significant decreases in energy metabolic enzymes and mitochondrial OXPHOS genes were identified, while cell adhesion and cytoskeleton molecules were upregulated (**Figure 12A, 12B**). Together, these alterations likely underlie the cell lineage differentiation defects. Similarly, these enteroid models showed a significant increase in Notch signaling molecules, which leads to the loss of tuft cell markers, such as *Pou2f3* and *Dclk1* (**Figure 12A**) (50). Supraphysiological activity of Notch signaling is associated with hyperproliferation of GI cancers (51). Consistent with this, our MVID patient-modeled pig (MYO5B(P663L)) enteroids demonstrated higher organoid forming rates and lower tuft cell numbers compared to wild type pig enteroids (**Figure 11C**). These observations suggest that MYO5B loss of function impacts cellular metabolism and Notch signaling of epithelial progenitors, leading to disruption of proper cell lineage differentiation.

As shown in our RNA-sequencing data of both tissue epithelial cells and enteroids, important chemokine ligands, such as *Ccl9* and *Ccl25*, were significantly downregulated (**Figure 13**). Enterocyte-produced CCL9 and CCL25 are required for the activation of gut-associated lymphoid tissues (GALT) that support epithelial barrier function (52, 53). GALT abnormalities have not been reported in MVID patients or animal models, but immune defects must be considered in long-term care. In future studies, the crosstalk between MYO5B deficient epithelial cells and mucosal immune system should be addressed.

### Therapeutic potential and limitation of LPAR5 activation in MVID

To support the impaired progenitor cell function by MYO5B loss, we have examined the effect of LPAR5 activation, which is required for intestinal stem cells (20). Treatment with the synthetic LPAR5 agonist, Compound-1, similarly improved apical expressions of NHE3 and SGLT1 in the MYO5BΔIEC and MYO5B(G519R) mice (**Figure 5D, 5E, 7A-D**). Apical NHE3 trafficking is ameliorated by natural lysophosphatidic acid (LPA)(18:1) in MYO5BΔIEC mice (12), and LPAR5 deficient mice indicate that NHE3 trafficking is mediated by LPAR5 in healthy epithelial cells (16, 17). Our present study suggests that LPAR5 agonists can recover NHE3 localization in MYO5B knockout as well as point mutant MVID models, bypassing a MYO5B-mediated trafficking pathway. Additionally, Compound-1 is a potential treatment for mislocalization of SGLT1, in which expression is limited to mature enterocytes (**Figure 7A-D**). Electrophysiological assays in the jejunum confirmed a significant improvement of SGLT1 activity by Compound-1 treatment in both MYO5BΔIEC and MYO5B(G519R) mice (**Figure 9B**). SGLT1 is water permeable, and loss-of-function mutations of SGLT1 cause congenital diarrhea referred to as glucose-galactose malabsorption (54–56). The partial improvement of functional brush border by Compound-1 treatment was further supported by the cellular distribution of the mitochondrial marker, NDUFB8 (**Figure 8**). In the treated MYO5BΔIEC and MYO5B(G519R) mouse tissues, NDUFB8 signals in the subapical zone of enterocytes was correlated with improved brush border structure. These observations suggest that ATP-dependent apical transporters and mitochondrial status were closely related and affected by LPAR5 agonist, Compound-1. LPAR5 activation may contribute to the functional maturation of absorptive enterocytes along with mitochondrial metabolism.

Despite these positive cellular effects, Compound-1 treatment had a limited effect on body weight loss only in MYO5B(G519R) male mice (**Figure 9A**). MYO5BΔIEC mice and MYO5B(G519R) female mice still lost 12–18% of original body weight within 4 days post tamoxifen. Although mechanisms underlying this sex- and genotype-specific effect of Compound-1 are still unclear, we speculate that non-epithelial cells are possibly activated to a different extent between males *vs*. females, and that the presence of mutant MYO5B(G519R) *vs*. absence of MYO5B differently affect epithelial cell responses to Compound-1-targeted non-epithelial cells. Vagal afferent nerves are expressing LPAR5 and likely activated by systemic Compound-1 administration resulting in reduced appetite. Additional nutrient supplementation and/or fluid therapy might be needed for total nutritional management.

Unlike the critical sodium transporters for water absorption, NHE3 and SGLT1, MUC13 localization was not remarkably improved by Compound-1 treatment (**Figure 10**). These results suggest that the intestinal brush border function is not completely re-established by LPAR5 activation alone. In wild type adult mice, Compound-1 was detected in plasma 1 hour after a bolus intraperitoneal injection (10 mg/kg) and disappeared (< 10 ng/ml) within 6 hours (data not shown). Compound-1 is designed as a metabolically stabilized LPA analogue, and its EC_50_ on human LPAR5 shows higher potency (0.26 nM) than natural LPA(18:1) (55 nM) (24). Further analysis of Compound-1 kinetics in human cells and identification of LPAR3 / LPAR5 expressing cell types in human tissues are required to develop clinical usage of this compound.

### Mechanism of Compound-1-mediated trophic effect

Daily Compound-1 treatment increased villus/crypt ratio by 150% in the small intestine of MYO5BΔIEC, but not in MYO5B(G519R) mice (**Figure 5C**), therefore, we have performed RNA-sequencing on MYO5BΔIEC enteroids with or without Compound-1. Compound-1 treatment of MYO5BΔIEC enteroids significantly (log2 fold change > |1|, adjusted *P* < 0.05) altered 85 genes compared to vehicle treatment. Out of these 85, 21 genes were differentially expressed in a comparison between MYO5BΔIEC and control enteroids (**Figure 12C**). Highlighted genes, which were MYO5B loss-increased and Compound-1-reversed, were implicated in cellular stress responses induced by inflammation or infection. This result indicates that the trophic effect of Compound-1 treatment seen *in vivo* was partly mediated by a direct effect on stressed epithelial cells. Additionally, Compound-1 significantly increased tuft cell differentiation *in vivo* in mouse models and in MVID pig enteroids (**Figure 11A, 11E**). These observations indicate an improvement of epithelial cell differentiation by LPAR5 activation in a cell autonomous pathway.

Recent mouse studies of LPAR5 deficient mice has revealed that LPAR5 both in epithelial cells and mucosal immune cells are essential for intestinal stem cell function (20). Intestinal progenitor apoptosis is acutely induced in total LPAR5 knockout mice, and stem cell self-renewal and organoid formation are inhibited by Lgr5+ cell-targeted LPAR5 loss, suggesting that LPAR5 activity is required for intestinal epithelial homeostasis. Intriguingly, decreased organoid forming capability of total LPAR5 knockout mice is rescued by supplementation with wild-type lymphocytes, implicating the presence of a reversible LPAR5 signaling pathway in intestinal stem cells (20). In addition to LPAR5, Compound-1 has an agonistic effect on human LPAR3 (24), which is diminished in the MYO5BΔIEC intestinal epithelial cells (**Figure 4D**). LPAR3 and LPAR5 are expressed in non-epithelial cells, such as in the immune and nervous systems (57, 58). Systemic treatment with Compound-1 may activate such non-epithelial cells which in turn may affect local epithelial cell function.

In conclusion, absence of MYO5B or a point mutation at MYO5B(G519R) disrupts intestinal progenitor cell metabolism and differentiation pathways. The LPAR5 agonist, Compound-1, ameliorates the defects of epithelial differentiation and functional maturation via epithelial cell-autonomous and non-epithelial cells in several MVID models. The present study provides further insights into unexplored roles of MYO5B specifically in intestinal stem cells and crosstalk between epithelial and mucosal immune cells.

## Acknowledgements

We acknowledge the Vanderbilt Chemical Synthesis Core for Compound-1 and UCM-05194, and the Translational Pathology Shared Resource supported by NCI/NIH Cancer Center support grant P30 CA68485-19. Special thanks to Dr. Michelle Reyzer in the Vanderbilt Mass Spectrometry Core for IMS data and to Dr. Evan Krystofiak in the Vanderbilt Cell Imaging Shared Resource for EM images. Graphical abstract was created with BioRender.com. This study was supported by the gift from Volpe Foundation and the National Institute of Health (NIH) RC2DK118640 and R01 DK48370 to JRG, and Vanderbilt Digestive Diseases Research Center Pilot and Feasibility Program (P30 DK058404), Vanderbilt GI SPORE Career Enhancement Award (P50 CA236733), and NIH R01 DK128190 to IK.

## Author Contributions

Conceptualization, I.K.; Methodology, J.T.R and I.K.; Software, M.E.B. and J.T.R.; Investigation, M.M., S.R., A.B., F.A., and I.K.; Data Curation, M.M., S.R. and F.A.; Resources, C.R. and J.R.G.; Writing – Original Draft, M.M., S.R., and I.K; Writing – Review & Editing, M.M., S.R., A.B., J.R.G., J.T.R. and I.K.; Funding Acquisition and Supervision, J.R.G. and I.K.

## References

1. Davidson GP, Cutz E, Hamilton JR, and Gall DG. Familial enteropathy: a syndrome of protracted diarrhea from birth, failure to thrive, and hypoplastic villus atrophy. Gastroenterology 75: 783–790, 1978.

2. Cutz E, Rhoads JM, Drumm B, Sherman PM, Durie PR, and Forstner GG. Microvillus inclusion disease: an inherited defect of brush-border assembly and differentiation. N Engl J Med 320: 646–651, 1989.

3. Erickson RP, Larson-Thome K, Valenzuela RK, Whitaker SE, and Shub MD. Navajo microvillous inclusion disease is due to a mutation in MYO5B. Am J Med Genet A 146a: 3117–3119, 2008.

4. Muller T, Hess MW, Schiefermeier N, Pfaller K, Ebner HL, Heinz-Erian P, Ponstingl H, Partsch J, Rollinghoff B, Kohler H, Berger T, Lenhartz H, Schlenck B, Houwen RJ, Taylor CJ, Zoller H, Lechner S, Goulet O, Utermann G, Ruemmele FM, Huber LA, and Janecke AR. MYO5B mutations cause microvillus inclusion disease and disrupt epithelial cell polarity. Nat Genet 40: 1163–1165, 2008.

5. Ruemmele FM, Schmitz J, and Goulet O. Microvillous inclusion disease (microvillous atrophy). Orphanet J Rare Dis 1: 22, 2006.

6. Thiagarajah JR, Kamin DS, Acra S, Goldsmith JD, Roland JT, Lencer WI, Muise AM, Goldenring JR, Avitzur Y, and Martin MG. Advances in Evaluation of Chronic Diarrhea in Infants. Gastroenterology 154: 2045–2059.e2046, 2018.

7. Burman A, Momoh M, Sampson L, Skelton J, Roland JT, Ramos C, Krystofiak E, Acra S, Goldenring JR, and Kaji I. Modeling of a Novel Patient-Based MYO5B Point Mutation Reveals Insights Into MVID Pathogenesis. Cell Mol Gastroenterol Hepatol 15: 1022–1026, 2023.

8. Kaji I, Thiagarajah JR, and Goldenring JR. Modeling the cell biology of monogenetic intestinal epithelial disorders. J Cell Biol 223: 2024.

9. Weis VG, Knowles BC, Choi E, Goldstein AE, Williams JA, Manning EH, Roland JT, Lapierre LA, and Goldenring JR. Loss of MYO5B in mice recapitulates Microvillus Inclusion Disease and reveals an apical trafficking pathway distinct to neonatal duodenum. Cell Mol Gastroenterol Hepatol 2: 131–157, 2016.

10. Engevik AC, Coutts AW, Kaji I, Rodriguez P, Ongaratto F, Saqui-Salces M, Medida RL, Meyer AR, Kolobova E, Engevik MA, Williams JA, Shub MD, Carlson DF, Melkamu T, and Goldenring JR. Editing Myosin VB Gene to Create Porcine Model of Microvillus Inclusion Disease, With Microvillus-Lined Inclusions and Alterations in Sodium Transporters. Gastroenterology 158: 2236–2249.e2239, 2020.

11. Engevik AC, Kaji I, Engevik MA, Meyer AR, Weis VG, Goldstein A, Hess MW, Muller T, Koepsell H, Dudeja PK, Tyska M, Huber LA, Shub MD, Ameen N, and Goldenring JR. Loss of MYO5B Leads to Reductions in Na(+) Absorption With Maintenance of CFTR-Dependent Cl(-) Secretion in Enterocytes. Gastroenterology 155: 1883–1897.e1810, 2018.

12. Kaji I, Roland JT, Watanabe M, Engevik AC, Goldstein AE, Hodges CA, and Goldenring JR. Lysophosphatidic Acid Increases Maturation of Brush Borders and SGLT1 Activity in MYO5B-deficient Mice, a Model of Microvillus Inclusion Disease. Gastroenterology 2020.

13. Kaji I, Roland JT, Rathan-Kumar S, Engevik AC, Burman A, Goldstein AE, Watanabe M, and Goldenring JR. Cell differentiation is disrupted by MYO5B loss through Wnt/Notch imbalance. JCI Insight 6: 2021.

14. Haber AL, Biton M, Rogel N, Herbst RH, Shekhar K, Smillie C, Burgin G, Delorey TM, Howitt MR, Katz Y, Tirosh I, Beyaz S, Dionne D, Zhang M, Raychowdhury R, Garrett WS, Rozenblatt-Rosen O, Shi HN, Yilmaz O, Xavier RJ, and Regev A. A single-cell survey of the small intestinal epithelium. Nature 551: 333–339, 2017.

15. Li C, Dandridge KS, Di A, Marrs KL, Harris EL, Roy K, Jackson JS, Makarova NV, Fujiwara Y, Farrar PL, Nelson DJ, Tigyi GJ, and Naren AP. Lysophosphatidic acid inhibits cholera toxin-induced secretory diarrhea through CFTR-dependent protein interactions. J Exp Med 202: 975–986, 2005.

16. Jenkin KA, He P, and Yun CC. Expression of lysophosphatidic acid receptor 5 is necessary for the regulation of intestinal Na(+)/H(+) exchanger 3 by lysophosphatidic acid in vivo. Am J Physiol Gastrointest Liver Physiol 315: G433–g442, 2018.

17. Lin S, Yeruva S, He P, Singh AK, Zhang H, Chen M, Lamprecht G, de Jonge HR, Tse M, Donowitz M, Hogema BM, Chun J, Seidler U, and Yun CC. Lysophosphatidic acid stimulates the intestinal brush border Na(+)/H(+) exchanger 3 and fluid absorption via LPA(5) and NHERF2. Gastroenterology 138: 649–658, 2010.

18. Konno T, Kotani T, Setiawan J, Nishigaito Y, Sawada N, Imada S, Saito Y, Murata Y, and Matozaki T. Role of lysophosphatidic acid in proliferation and differentiation of intestinal epithelial cells. PLoS One 14: e0215255, 2019.

19. Liang Z, and Yun CC. Compensatory Upregulation of LPA(2) and Activation of the PI3K-Akt Pathway Prevent LPA(5)-Dependent Loss of Intestinal Epithelial Cells in Intestinal Organoids. Cells 11: 2022.

20. Liang Z, He P, Han Y, and Yun CC. Survival of Stem Cells and Progenitors in the Intestine Is Regulated by LPA_5_-Dependent Signaling. Cellular and Molecular Gastroenterology and Hepatology 14: 129–150, 2022.

21. Liu W, Hopkins AM, and Hou J. The development of modulators for lysophosphatidic acid receptors: A comprehensive review. Bioorg Chem 117: 105386, 2021.

22. Meduri B, Pujar GV, Durai Ananda Kumar T, Akshatha HS, Sethu AK, Singh M, Kanagarla A, and Mathew B. Lysophosphatidic acid (LPA) receptor modulators: Structural features and recent development. Eur J Med Chem 222: 113574, 2021.

23. González-Gil I, Zian D, Vázquez-Villa H, Hernández-Torres G, Martínez RF, Khiar-Fernández N, Rivera R, Kihara Y, Devesa I, Mathivanan S, Del Valle CR, Zambrana-Infantes E, Puigdomenech M, Cincilla G, Sanchez-Martinez M, Rodríguez de Fonseca F, Ferrer-Montiel AV, Chun J, López-Vales R, López-Rodríguez ML, and Ortega-Gutiérrez S. A Novel Agonist of the Type 1 Lysophosphatidic Acid Receptor (LPA(1)), UCM-05194, Shows Efficacy in Neuropathic Pain Amelioration. J Med Chem 63: 2372–2390, 2020.

24. Jiang G, Inoue A, Aoki J, and Prestwich GD. Phosphorothioate analogs of sn-2 radyl lysophosphatidic acid (LPA): metabolically stabilized LPA receptor agonists. Bioorg Med Chem Lett 23: 1865–1869, 2013.

25. Berg S, Kutra D, Kroeger T, Straehle CN, Kausler BX, Haubold C, Schiegg M, Ales J, Beier T, Rudy M, Eren K, Cervantes JI, Xu B, Beuttenmueller F, Wolny A, Zhang C, Koethe U, Hamprecht FA, and Kreshuk A. ilastik: interactive machine learning for (bio)image analysis. Nat Methods 16: 1226–1232, 2019.

26. The MathWorks Inc. MATLAB version: 9.13.0 (R2022b): https://www.mathworks.com.

27. Kaji I, Roland JT, Watanabe M, Engevik AC, Goldstein AE, Hodges CA, and Goldenring JR. Lysophosphatidic Acid Increases Maturation of Brush Borders and SGLT1 Activity in MYO5B-deficient Mice, a Model of Microvillus Inclusion Disease. Gastroenterology 159: 1390–1405.e1320, 2020.

28. Mahe MM, Aihara E, Schumacher MA, Zavros Y, Montrose MH, Helmrath MA, Sato T, and Shroyer NF. Establishment of Gastrointestinal Epithelial Organoids. Curr Protoc Mouse Biol 3: 217–240, 2013.

29. Mortazavi A, Williams BA, McCue K, Schaeffer L, and Wold B. Mapping and quantifying mammalian transcriptomes by RNA-Seq. Nat Methods 5: 621–628, 2008.

30. Babicki S, Arndt D, Marcu A, Liang Y, Grant JR, Maciejewski A, and Wishart DS. Heatmapper: web-enabled heat mapping for all. Nucleic Acids Res 44: W147–153, 2016.

31. Moriyama R, and Fukushima N. Expression of lysophosphatidic acid receptor 1 in the adult female mouse pituitary gland. Neurosci Lett 741: 135506, 2021.

32. Huh WJ, Roland JT, Asai M, and Kaji I. Distribution of duodenal tuft cells is altered in pediatric patients with acute and chronic enteropathy. Biomed Res 41: 113–118, 2020.

33. Ling C, Versloot CJ, Arvidsson Kvissberg ME, Hu G, Swain N, Horcas-Nieto JM, Miraglia E, Thind MK, Farooqui A, Gerding A, van Eunen K, Koster MH, Kloosterhuis NJ, Chi L, ChenMi Y, Langelaar-Makkinje M, Bourdon C, Swann J, Smit M, de Bruin A, Youssef SA, Feenstra M, van Dijk TH, Thedieck K, Jonker JW, Kim PK, Bakker BM, and Bandsma RHJ. Rebalancing of mitochondrial homeostasis through an NAD(+)-SIRT1 pathway preserves intestinal barrier function in severe malnutrition. EBioMedicine 96: 104809, 2023.

34. Moschandrea C, Kondylis V, Evangelakos I, Herholz M, Schneider F, Schmidt C, Yang M, Ehret S, Heine M, Jaeckstein MY, Szczepanowska K, Schwarzer R, Baumann L, Bock T, Nikitopoulou E, Brodesser S, Krüger M, Frezza C, Heeren J, Trifunovic A, and Pasparakis M. Mitochondrial dysfunction abrogates dietary lipid processing in enterocytes. Nature 625: 385–392, 2024.

35. Berger E, Rath E, Yuan D, Waldschmitt N, Khaloian S, Allgäuer M, Staszewski O, Lobner EM, Schöttl T, Giesbertz P, Coleman OI, Prinz M, Weber A, Gerhard M, Klingenspor M, Janssen KP, Heikenwalder M, and Haller D. Mitochondrial function controls intestinal epithelial stemness and proliferation. Nat Commun 7: 13171, 2016.

36. Rath E, Moschetta A, and Haller D. Mitochondrial function - gatekeeper of intestinal epithelial cell homeostasis. Nat Rev Gastroenterol Hepatol 15: 497–516, 2018.

37. Smith AL, Whitehall JC, Bradshaw C, Gay D, Robertson F, Blain AP, Hudson G, Pyle A, Houghton D, Hunt M, Sampson JN, Stamp C, Mallett G, Amarnath S, Leslie J, Oakley F, Wilson L, Baker A, Russell OM, Johnson R, Richardson CA, Gupta B, McCallum I, McDonald SA, Kelly S, Mathers JC, Heer R, Taylor RW, Perkins ND, Turnbull DM, Sansom OJ, and Greaves LC. Age-associated mitochondrial DNA mutations cause metabolic remodelling that contributes to accelerated intestinal tumorigenesis. Nat Cancer 1: 976–989, 2020.

38. Cheng CW, Biton M, Haber AL, Gunduz N, Eng G, Gaynor LT, Tripathi S, Calibasi-Kocal G, Rickelt S, Butty VL, Moreno-Serrano M, Iqbal AM, Bauer-Rowe KE, Imada S, Ulutas MS, Mylonas C, Whary MT, Levine SS, Basbinar Y, Hynes RO, Mino-Kenudson M, Deshpande V, Boyer LA, Fox JG, Terranova C, Rai K, Piwnica-Worms H, Mihaylova MM, Regev A, and Yilmaz OH. Ketone Body Signaling Mediates Intestinal Stem Cell Homeostasis and Adaptation to Diet. Cell 178: 1115–1131.e1115, 2019.

39. Chen L, Vasoya RP, Toke NH, Parthasarathy A, Luo S, Chiles E, Flores J, Gao N, Bonder EM, Su X, and Verzi MP. HNF4 Regulates Fatty Acid Oxidation and Is Required for Renewal of Intestinal Stem Cells in Mice. Gastroenterology S0016-5085(0019)41587–41584, 2019.

40. Stine RR, Sakers AP, TeSlaa T, Kissig M, Stine ZE, Kwon CW, Cheng L, Lim H-W, Kaestner KH, Rabinowitz JD, and Seale P. PRDM16 Maintains Homeostasis of the Intestinal Epithelium by Controlling Region-Specific Metabolism. Cell stem cell 25: 830–845.e838, 2019.

41. Sheng YH, Lourie R, Linden SK, Jeffery PL, Roche D, Tran TV, Png CW, Waterhouse N, Sutton P, Florin TH, and McGuckin MA. The MUC13 cell-surface mucin protects against intestinal inflammation by inhibiting epithelial cell apoptosis. Gut 60: 1661–1670, 2011.

42. Rathan-Kumar S, Roland JT, Momoh M, Goldstein A, Lapierre LA, Manning E, Mitchell L, Norman J, Kaji I, and Goldenring JR. Rab11FIP1-deficient mice develop spontaneous inflammation and show increased susceptibility to colon damage. Am J Physiol Gastrointest Liver Physiol 323: G239–G254, 2022.

43. McKinley ET, Sui Y, Al-Kofahi Y, Millis BA, Tyska MJ, Roland JT, Santamaria-Pang A, Ohland CL, Jobin C, Franklin JL, Lau KS, Gerdes MJ, and Coffey RJ. Optimized multiplex immunofluorescence single-cell analysis reveals tuft cell heterogeneity. JCI Insight 2: 2017.

44. Gerbe F, Sidot E, Smyth DJ, Ohmoto M, Matsumoto I, Dardalhon V, Cesses P, Garnier L, Pouzolles M, Brulin B, Bruschi M, Harcus Y, Zimmermann VS, Taylor N, Maizels RM, and Jay P. Intestinal epithelial tuft cells initiate type 2 mucosal immunity to helminth parasites. Nature 529: 226–230, 2016.

45. Burman A, and Kaji I. Luminal Chemosensory Cells in the Small Intestine. Nutrients 13: 2021.

46. Vogel GF, Janecke AR, Krainer IM, Gutleben K, Witting B, Mitton SG, Mansour S, Ballauff A, Roland JT, Engevik AC, Cutz E, Muller T, Goldenring JR, Huber LA, and Hess MW. Abnormal Rab11-Rab8-vesicles cluster in enterocytes of patients with microvillus inclusion disease. Traffic 18: 453–464, 2017.

47. Roland JT, Bryant DM, Datta A, Itzen A, Mostov KE, and Goldenring JR. Rab GTPase-Myo5B complexes control membrane recycling and epithelial polarization. Proc Natl Acad Sci U S A 108: 2789–2794, 2011.

48. Puthalakath H, Villunger A, O’Reilly LA, Beaumont JG, Coultas L, Cheney RE, Huang DC, and Strasser A. Bmf: a proapoptotic BH3-only protein regulated by interaction with the myosin V actin motor complex, activated by anoikis. Science 293: 1829–1832, 2001.

49. Hausmann M, Leucht K, Ploner C, Kiessling S, Villunger A, Becker H, Hofmann C, Falk W, Krebs M, Kellermeier S, Fried M, Schölmerich J, Obermeier F, and Rogler G. BCL-2 modifying factor (BMF) is a central regulator of anoikis in human intestinal epithelial cells. J Biol Chem 286: 26533–26540, 2011.

50. VanDussen KL, Carulli AJ, Keeley TM, Patel SR, Puthoff BJ, Magness ST, Tran IT, Maillard I, Siebel C, Kolterud A, Grosse AS, Gumucio DL, Ernst SA, Tsai YH, Dempsey PJ, and Samuelson LC. Notch signaling modulates proliferation and differentiation of intestinal crypt base columnar stem cells. Development 139: 488–497, 2012.

51. Demitrack ES, and Samuelson LC. Notch regulation of gastrointestinal stem cells. J Physiol 594: 4791–4803, 2016.

52. Zhao X, Sato A, Dela Cruz CS, Linehan M, Luegering A, Kucharzik T, Shirakawa AK, Marquez G, Farber JM, Williams I, and Iwasaki A. CCL9 is secreted by the follicle-associated epithelium and recruits dome region Peyer’s patch CD11b+ dendritic cells. J Immunol 171: 2797–2803, 2003.

53. Campbell DJ, and Butcher EC. Intestinal attraction: CCL25 functions in effector lymphocyte recruitment to the small intestine. J Clin Invest 110: 1079–1081, 2002.

54. Martin MG, Turk E, Lostao MP, Kerner C, and Wright EM. Defects in Na+/glucose cotransporter (SGLT1) trafficking and function cause glucose-galactose malabsorption. Nat Genet 12: 216–220, 1996.

55. Wright EM, Loo DD, and Hirayama BA. Biology of human sodium glucose transporters. Physiol Rev 91: 733–794, 2011.

56. Han L, Qu Q, Aydin D, Panova O, Robertson MJ, Xu Y, Dror RO, Skiniotis G, and Feng L. Structure and mechanism of the SGLT family of glucose transporters. Nature 601: 274–279, 2022.

57. Yung YC, Stoddard NC, Mirendil H, and Chun J. Lysophosphatidic Acid signaling in the nervous system. Neuron 85: 669–682, 2015.

58. Dacheux MA, Norman DD, Tigyi GJ, and Lee SC. Emerging roles of lysophosphatidic acid receptor subtype 5 (LPAR5) in inflammatory diseases and cancer. Pharmacol Ther 245: 108414, 2023.

