## Supplementary Figure 1 for "Altered cellular metabolic pathway and epithelial cell maturation induced by MYO5B defects are partially reversible by LPAR5 activation"

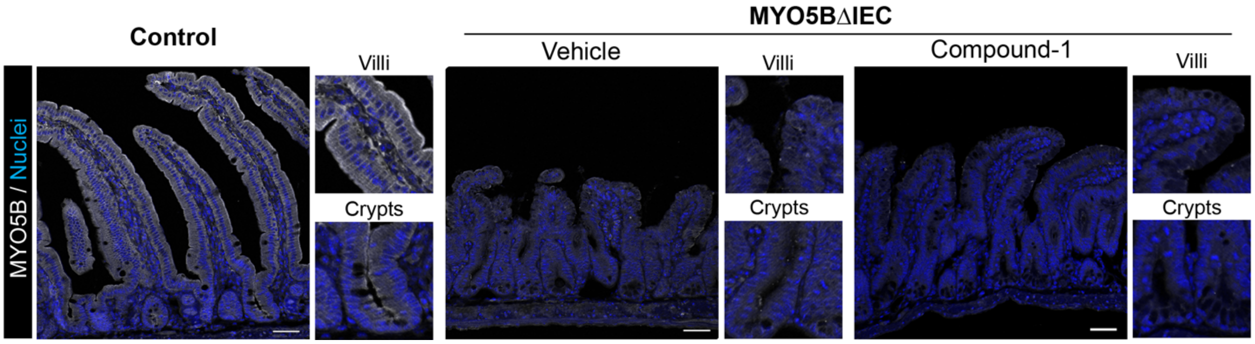


**Supplementary Figure 1.** Absence of MYO5B expression in MYO5BΔIEC mouse intestines following tamoxifen injection with or without Compound-1 treatment. Immunostaining for MYO5B (1:500, NBP1-87746, Novus) was performed on intestinal sections of control and MYO5BΔIEC mice. In control tissues, all epithelial cells along crypt-villous axis express MYO5B with intense localization in the subapical area. MYO5B signals are abolished in most epithelial cells in MYO5BΔIEC intestines with vehicle or Compound-1 treatment.
